# Septins Enable T Cell Contact Guidance *via* Amoeboid-Mesenchymal Switch

**DOI:** 10.1101/2023.09.26.559597

**Authors:** Alexander S. Zhovmer, Alexis Manning, Chynna Smith, Jian Wang, Xuefei Ma, Denis Tsygankov, Nikolay V. Dokholyan, Alexander X. Cartagena-Rivera, Rakesh K. Singh, Erdem D. Tabdanov

**Affiliations:** Center for Biologics Evaluation & Research, U.S. Food and Drug Administration, Silver Spring, MD, USA; Section on Mechanobiology, National Institute of Biomedical Imaging and Bioengineering, National Institutes of Health, Bethesda, MD, USA; Departments of Pharmacology, Penn State College of Medicine, The Pennsylvania State University, Hershey, PA, USA; Wallace H. Coulter Department of Biomedical Engineering, Georgia Institute of Technology and Emory University, Atlanta, GA, USA; Department of Biochemistry and Molecular Biology, Penn State College of Medicine, The Pennsylvania State University Hershey-Hummelstown, PA, USA; Department of Obstetrics & Gynecology, University of Rochester Medical Center, Rochester, NY, USA; Penn State Cancer Institute, Penn State College of Medicine, The Pennsylvania State University, Hershey, PA, USA

**Keywords:** guidance, lymphocyte, septin, dynein, migration

## Abstract

Lymphocytes exit circulation and enter in-tissue guided migration toward sites of tissue pathologies, damage, infection, or inflammation. By continuously sensing and adapting to the guiding chemo-mechano-structural properties of the tissues, lymphocytes dynamically alternate and combine their amoeboid (non-adhesive) and mesenchymal (adhesive) migration modes. However, which mechanisms guide and balance different migration modes are largely unclear. Here we report that suppression of septins GTPase activity induces an abrupt amoeboid-to-mesenchymal transition of T cell migration mode, characterized by a distinct, highly deformable integrin-dependent immune cell contact guidance. Surprisingly, the T cell actomyosin cortex contractility becomes diminished, dispensable and antagonistic to mesenchymal-like migration mode. Instead, mesenchymal-like T cells rely on microtubule stabilization and their non-canonical dynein motor activity for high fidelity contact guidance. Our results establish septin’s GTPase activity as an important on/off switch for integrin-dependent migration of T lymphocytes, enabling their dynein-driven fluid-like mesenchymal propulsion along the complex adhesion cues.

**SIGNIFICANCE STATEMENT:** Deciphering mechanisms of guided lymphocyte migration paves the way towards effective immunotherapies for the extracellular matrix-rich tissues, such as solid tumors. Here we demonstrate that T cell septins’ GTPase activity regulates both actomyosin and microtubules, alternately enhancing either of these two major motor systems. Surprisingly, the suppression of septin GTPase activity also induces a highly guided integrin-dependent mesenchymal-like migration directed by the extracellular matrix proteins. The phenomenon of guided mesenchymal-like migration of T cells relies on the microtubules and microtubule-based dynein motors that are responsible for the force generation, powering guided T cell motility. This finding opens a new perspective for future studies of septin GTPases in a context of the optimisation of T cell-based immunotherapies for the solid tissues.

## INTRODUCTION

Surveilling immune cells experience extreme morphological deformations during migration throughout a variety of confining and structurally interactive tissue microenvironments (1). Inadequate migration of immune cells in tissues contributes to the formation of immuno-resistant cancer sanctuaries and autoimmune, allergic, and neurodegenerative disorders (2–5). For adapting and tuning their motility within a specific encountered microenvironment, lymphocytes sense guiding cues of tissue microenvironments and adaptively remodel their cytoskeletal organization for combining or alternating between appropriate migration modes (6–8). For instance, a non-adhesive amoeboid (9, 10) and adhesive mesenchymal-like migration modes (8, 11) are combined into a ‘random-walk’ motility (6, 8), which additionally may be accompanied by a 3D ‘paddling’ in fluids (12).

The elements of adhesive (*i.e.*, integrin-dependent mesenchymal-like) migration are proposed to play an important role in the directional propulsion of amoeboid T cells within a structurally discrete tissue microenvironment (7) or during transmigration (13). For example, cascades of integrin-dependent phenotype switching events and corresponding cytoskeletal reorganizations guide the recruitment of lymphocytes to tissues during the directional transmigration of immune cells between the bloodstream and organs (14, 15). Multiple direct observations in various experiments suggest that the cytoskeletal organization of transmigrating immune cells differs from that of amoeboid immune cells (16–18), including the activation of signaling pathways specific for mesenchymal-like migration (16, 19–22).

However, despite the significant clinical and scientific interest, it is still largely unknown how immune cells adapt their cytoskeleton for optimized motility. In part, these mechanisms are difficult to decipher due to the extreme structural and signaling entanglement between different cytoskeleton components. For example, studies demonstrate that the microtubular component, which is prevalent during mesenchymal-like migration, mechanically integrates into the F-actin cytoskeleton, which in turn is prevalent during amoeboid migration, *via* force-generating dyneins (23, 24), forming a singular microtubule-actomyosin force transmission system, hence, mechanically converging to the common cell adhesion receptors *(e.g.*, integrins). Furthermore, both actomyosin and microtubule components crosstalk with septin GTP-binding proteins, which, similarly to actomyosin and microtubules, can regulate the motility of immune cells (9).

Historically, septins are known to reside within the actomyosin cortex, participating in microtubule abscission and microtubule-driven cytokinesis (25, 26). For example, a septin-7 isoform is reported as the MAP4 binding factor that prevents MAP4 from binding to microtubules, thus suppressing MAP4-mediated microtubule stabilization (27, 28). On the contrary, septin-9’s direct binding to microtubules causes suppression of microtubules plus-end instability, leading to the enhancement of the microtubular apparatus (29), perhaps *via* recently found MAP-like motif in SEPT9_i1 isoform (30), and suggesting a more complex role for septins, which, at the minimum, may act both as actomyosin components and as bi-directional regulators of microtubule-associated signaling. Notably, loss of septin-9 (SEPT9^-/-^) leads to a partial loss of fibroblasts, *i.e.*, mesenchymal cells, transmigration efficiency, highlighting a possible causative link between mesenchymal-like T cell transmigration, septins activity (31), and microtubule system (32, 33).

These studies highlight the significant mechanobiological crosstalk between actomyosin, microtubule, and septin cytoskeletal components. However, the role of this crosstalk in the guided migration of immune cells remains unexplored. Here we show that modulating septins GTPase activity in T lymphocytes provides switching towards directional migration upon sensing of guiding microenvironment cues. In particular, we show that inhibition of septins’ GTPase activity in T cells causes an abrupt loss of amoeboid migration and transition towards mesenchymal-like motility characterized by distinct contact guidance and stabilization of microtubules *via* MAP4 and HDAC6 signaling.

## RESULTS

### Testing of novel septins GTPase inhibitor in mesenchymal cells

To study the role of septin GTPase activity during the amoeboid-to-mesenchymal transition in mammalian immune cells, we use the UR214-9, which is a non-toxic fluoride-derived analog of forchlorfenuron (FCF) - the inhibitor of the fungal (*i.e.,* yeast) septin GTPase (34)^,(35–37)^. For that, we first characterize septin-docking energies for the UR214-9 molecule compared to GTP, GDP, and FCF at the septin molecules’ GTPase sites. Specifically, we aim for an *in silico* molecular modeling for septin-inhibitor and septin-GTP/GDP docking and energies of interactions. Results demonstrate the competitive and comparable interaction energies for all listed molecules, while UR214-9 also displays more optimal docking energies than that of FCF for human septin-2, -6, -7, and -9 **(Figure 1a)**. More detailed visualization of the *in silico*-simulated UR214-9, GTP, GDP, and FCF dockings into pocket 1 of the human septin-7 are shown in **Figure 1b-1**, and dockings into pocket 2 of the human septin 7 are shown in **Figure 1b-2**. The complete sets of the molecular docking configurations for all filamentous septins, *i.e.*, septin-2, -6, -7, and -9, are shown in **Figure SI1a**, while the corresponding calculated *in silico* energies of the docking interactions within their pockets 1 and 2 are shown in **Figure SI1b**.

**Figure 1.**
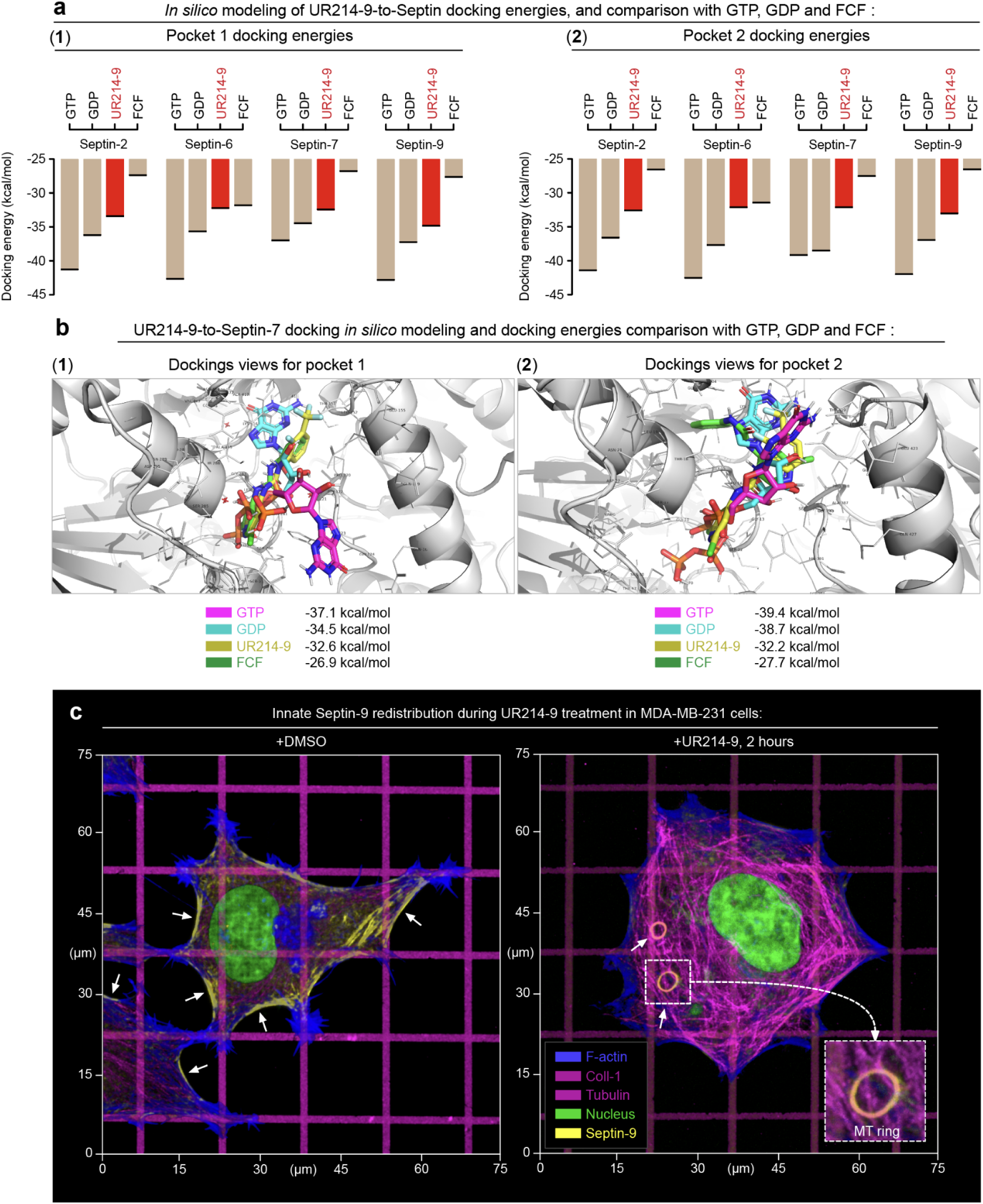
*In silico* simulations of molecular docking for the key filaments-related septin isoforms display a competitive UR214-9-septins interactions energies in comparison to GTP, GDP and FCF. **(a)** *In silico*-computationally derived energies of GTP, GDP, UR214-9, and FCF docking to pocket-1 and -2 of Septin-7 isoform. Note, that UR214-9-septin docking energy is more optimal, than that of FCF across all key filaments-related septin isoforms. **(b)** Examples of *in silico*-simulated 3D models of molecular docking of GTP, GFP, UR214-9 and FCF to the septin-7’s pockets 1 and 2. Additional models for Septin-2, -6, and -9 are included in **Supplemental Figure SI1**. **(c)** In control conditions **(+DMSO)**, biaxial spreading of mesenchymal MDA-MB231 cells along the orthogonal collagen-1 grid results in the polygonal cell architecture. Free cell edges feature non-compensated linear anisotropy along the arcs, resulting in the enhancement of linear contractile actomyosin filaments. Consequently, free cell edges feature the highest tension and develop the most of the stress-fibers. The tensile actomyosin (stress-fibers) accumulates septin-9 (*arrows*), while acute septin inhibition **(+UR214-9)** induces septin-9 translocation from actomyosin onto the microtubules, where they induce dynein-dependent microtubules circularization into the rings (*arrows*). Inset in a dashed box shows the detailed SEP9-decorated MT rings.

**Figure 2.**
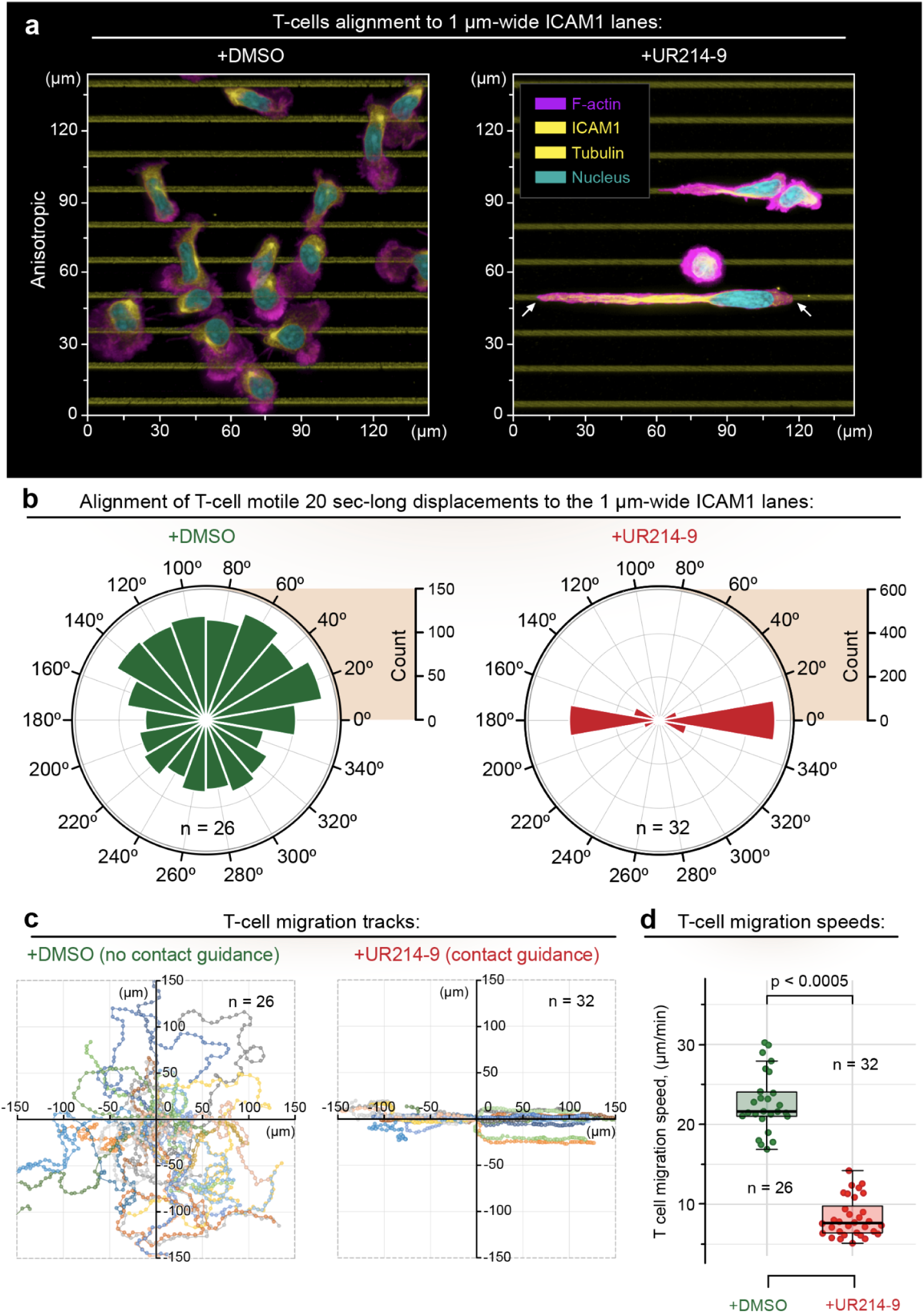
Acute inactivation of septins GTPase activity in CD4+ T lymphocytes activates amoeboid-to-mesenchymal-like phenotypic transitioning, characterized by distinct contact guidance along the adhesive ICAM-1-cues and slow mesenchymal-like migration. **(a)** hCD4+ T cells in control conditions **(+DMSO)** maintain amoeboid phenotype, featuring neither spreading nor alignment along the adhesion cues (ICAM1 anisotropic parallel lines, see **Movies 1** and **2 (before UR214-9)**). Septin inhibition **(+UR214-9)** induces phenotype transitioning towards mesenchymal-like behavior, which results in the prominent cell adhesion, alignment, and spreading along the ICAM1 lines **(Movies 1** and **2 (after UR214-9), Movie 3)**. Comparative analysis of migratory dynamics in control **(+DMSO)** and septin-inhibited **(+UR214-9)** T cells for: **(b)** migration directionality, **(c)** individual cell migration tracks, and **(d)** migration speeds, indicates that septins inhibition leads to the transition towards extreme, mesenchymal-like migration and alignment to the contact guidance cues, characterized by high fidelity of the T cell migration tracks to the ICAM1 lines architecture, and accompanied with the significant loss of T cell migration speed, inherent to the slow mesenchymal migratory dynamics.

To confirm that UR214-9 acts like FCF, *i.e.*, causes loss of septin-actomyosin interactions, we test septin-inhibitory effects of UR214-9 using adherent MDA-MB-231 cells, the human breast adenocarcinoma, which unlike T cells develop easily distinguishable actomyosin structures, *i.e.*, stress-fibers, that are stabilized by septins (38). To reinforce the buildup of visible tensile stress-fibers in MDA-MB-231 cells, we print collagen type-1 orthogonal grid with 1-µm-wide micro-lanes, spaced with 15 µm pitch on rigid polyacrylamide gel (shear modulus G’=50 kPa), to enhance the mechanical and spatial anisotropy of tensile actomyosin. The ‘fibrous’ biomimetic surrogate of the extracellular matrix induces biaxial polygonal spreading of mesenchymal cells (37, 39), with cell configurations featuring linearly stressed arching free cell edges **(Figure 1c)**, where the actomyosin filaments experience the highest linear mechanical tension (40, 41). Thus, the actomyosin linear tension in the cell’s arcs subsequently reinforces the buildup of tensile stress fibers *via* positive mechanosensory feedback **(Figure 1c**, **+DMSO**, *arrows*) (39, 41). We note that septin distribution in the polygonal MDA-MB-231 cells shows that septin-9 replicates spatial distribution of the tensile actomyosin (*i.e.*, stress fibers) with septin filaments showing strong spatial and structural affinity towards the arching free cell edges **(Figure 1c**, **+DMSO,** single cell view, *arrows*; **Figure SI2**, **+DMSO**, population-wide view, *arrows*). Inhibition of septins GTPase activity with UR214-9 abrogates the accumulation of septin-9 within free cell edges and drives septins translocation onto the microtubules **(Figure 1c**, **+UR214-9**, single cell view, *arrows*; **Figure SI2**, **+UR214-9**, population-wide view, *arrows*).

We observe that translocation of septin-9 onto the microtubules in the fully-spread polygonal MDA-MB-231 cells increases microtubule density **(Figure 1c**, **+DMSO** *vs.* **+UR214-9)** and induces microtubule curling into the rings in ∼80% of the cell population **(Figure SI2**, **+UR214-9**, *arrows*), complementing our previous report on septin-9-mediated microtubule reorganization in mesenchymal cells (37). Potentially, the accumulation of GTPase-inactive septin-9 on microtubules may occur either *via* shuttling between: (i) septins associated with tensile septin-crosslinked actomyosin cytoskeleton, (ii) septins deposited on microtubules, and (iii) cytosolic septin-9, which adds to the 7-6-2-2-6-7 septin hexamers resulting in the formation of the 9-7-6-2-2-6-7-9 octamers (26, 42, 43), or directly, *via* a recently identified MAP-like motif in the SEPT9_i1 (*i.e.* septin-9, isoform 1), which facilitates SEPT9_i1 direct binding to the microtubules (30).

### Septins inhibition activates mesenchymal-like contact guidance in T cells

Enhancement of the microtubules density due to septins translocation onto the microtubules *via* UR214-9-induced loss of septins association with the actomyosin in MDA-MB-231 cells may replicate traits specific for epithelial-mesenchymal transition (EMT) (44) and mesenchymal cell phenotypes (45). For example, inhibition of actomyosin contractility in MDA-MB-231 (46), MDA-MB-468 (39), and human epithelial cells (47) routinely leads to mesenchymal-like enhancement of cell morphological conformity to the adhesion cues. Therefore, we hypothesize that inhibition of septin GTPase activity may also enhance mesenchymal-like T cell conformity to 2D guidance cues, explaining the observed instances of *in vivo* migratory compliance of T cells to the adhesion guidance cues (48).

To test our hypothesis, we put human CD4+ T cells on a substrate designed to reveal cell shape and migration compliance in the form of their morphological and migratory alignment to the 2D anisotropic adhesive contact guidance cues, *i.e.,* 1-µm-wide, 15-µm-pitched ICAM1 micro-lanes **(Figure 2a)**. As reported previously (8), in the control conditions **(+DMSO)**, T cells show amoeboid phenotype with typical meandering ‘random-walk’ crawling and display no detectable structural or migratory alignment to the ICAM1 micro-lanes **(Figure 2a-c**, **+DMSO**, **Movies 1** and **2)**. The detailed analysis of T cell migration trajectories on the ICAM1 micro-lanes shows that T cells display a random distribution of directionalities, measured within 20-second-long timeframes **(Figure 2b**, **+DMSO)**, and highly meandering migration tracks of individual cells **(Figure 2c**, **+DMSO)**.

Upon acute pharmacological inhibition of septin GTPase activity **(+UR214-9**, t≪1 hour), amoeboid T cells abruptly transition towards a mesenchymal-like morphological alignment and elongation along the 1-µm-wide ICAM1 lanes **(Figure 2a**, **+UR214-9**, t = 30 min; **Movies 1** and **2**, **after UR214-9**, **Movie 3)**, which is also accompanied by a highly aligned and persistent migration along the ICAM1 micro-lanes **(Figure 2b-c**, **+UR214-9**, **Movie 3)**. Notably, while amoeboid T cells migrate at highly dynamic ∼22 µm/min velocity **(Figure 2d**, **+DMSO**), the mesenchymal-like T cells migrate at more conventional ∼7.5 µm/min cell speed (**Figure 2d**, **+UR214-9**), which may reflect an overall higher dynamics of the non-adhesive amoeboid T cell cortex, compared to the cytoskeletal reorganization configured by the integrin-driven mesenchymal-like adhesion and migration mechanisms (6).

While reprogramming of T cells towards mesenchymal-like motility manifests as directional alignment (*i.e.,* contact guidance) of T cells along the adhesive ICAM1 micro-lanes **(Figure 2a**,**+DMSO** *vs.* **+UR214-9)**, increasing the geometric complexity of the ICAM1 adhesion cues from the parallel lanes to the orthogonal grids also leads to the equal enhancement of the T cell’s morphological congruence to the underlying adhesion geometry **(Figure 3a**, **+DMSO** *vs.* **+UR214-9)**, quantifiable as T cell morphological complexity **(Figure SI3)**. Thus, T cell alignment to the ICAM1 geometry cannot be explained simply by an overall loss of the cytoskeletal organization complexity, such as linear polarization of the T cell cytoskeleton. Instead, it reflects an elevated T cell sensitivity and congruence towards the adhesion geometry, *i.e.*, contact guidance. Thus, a question arises on which mechanisms facilitate an enhanced T cell’s conformity to the adhesion cues.

**Figure 3.**
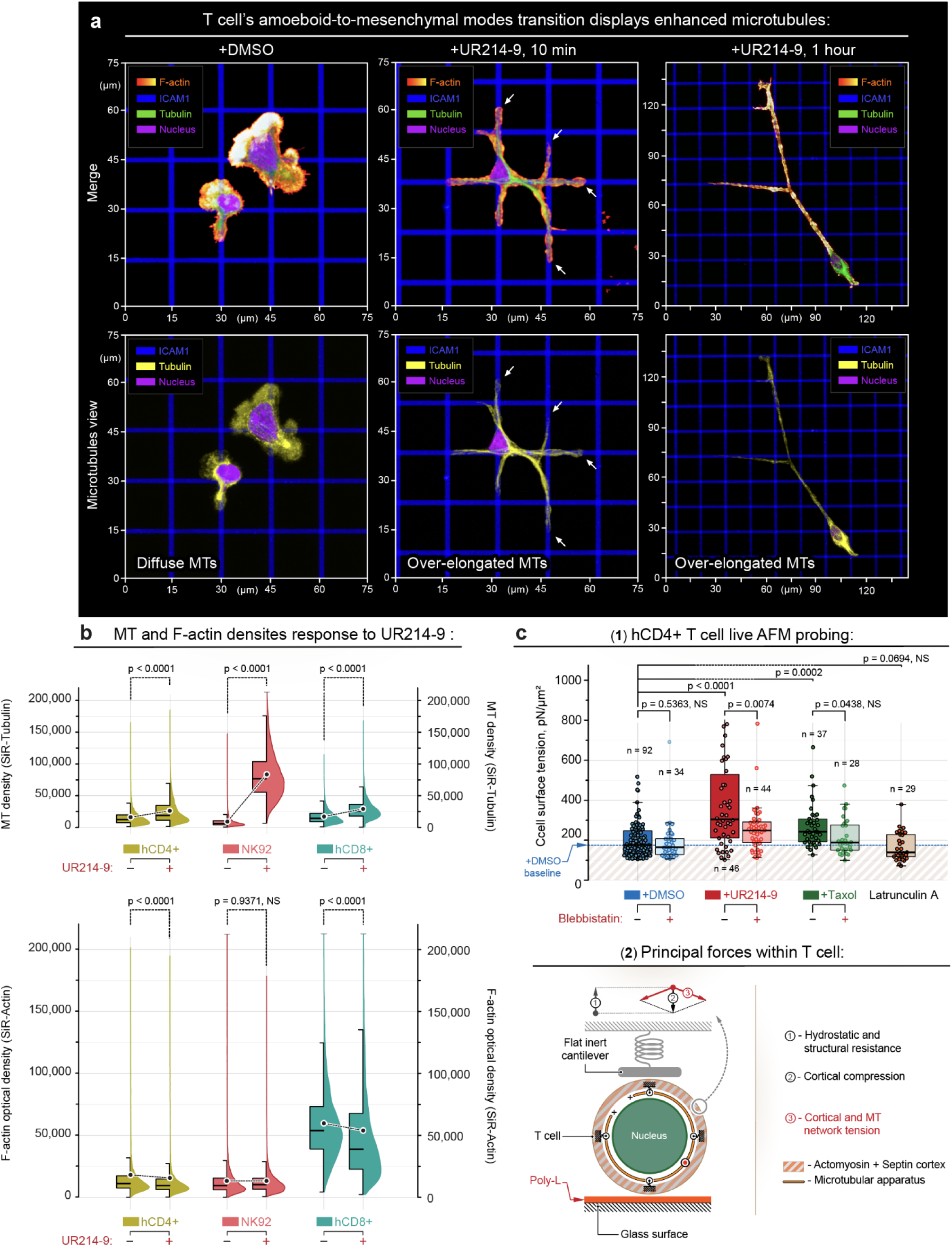
Inhibition of septin’s GTPase activity increases microtubules density and T cell rigidity, while reducing F-actin density. **(a)** Visualization of T cell (hCD4+) amoeboid-to-mesenchymal-like phenotypic transitioning. Amoeboid T cells are characterized by their morphological unresponsiveness to the geometry of the underlying adhesive ICAM1 grid **(+DMSO**, **Movie 4).** Upon inhibition of septins, they acutely switch towards mesenchymal-like morphological congruency and form multiple dendritic-like protrusions along the ICAM1 grid geometry **(+UR214-9**, 10 min, **Movie 5)**. Additionally, prolonged septin inhibition **(+UR214-9**, 1 hour) causes T cell over-stretching in a mesenchymal-like manner, with T cell adhesion and spreading along the ICAM1 adhesion grid. *Top* - Visualization of the F-actin, nucleus and microtubules. *Bottom* - Separate visualization of microtubules’ density and length: Short microtubule filaments and diffuse non-filamentous tubulin **(+DMSO)** transform into the over-elongated and dense microtubules (*arrows*) during septins inhibition **(+UR214-9**, 10 min), further evolving into the extremely elongated microtubules **(+UR214-9**, t≥1 hour). **(b)** Live lymphocytes flow cytometry analysis of microtubule (SiR-Tubulin) and filamentous (F)-actin (SiR-Actin) density changes in the hCD4+ T-, NK92, and hCD8+ T-lymphocytes in response to the acute septin inhibition with UR214-9. *Top* - Microtubule density in control and UR214-9-treated cells, as measured in live cells by staining with the microtubule-specific SiR-Tubulin (1 hour staining time). *Bottom* - F-actin density in control and UR214-9-treated cells, as measured in live cells by F-actin-specific labeling with SiR-actin (1 hour staining time). Microtubule density increases in response to UR214-9 treatment in hCD4+, NK92 and hCD8+ lymphocytes **(+UR214-9**, t≥1 hour), showing ∼60% (hCD4+), ∼1000% (NK92) or ∼90% (hCD8+) increase of microtubules density, *per cell*, compared to the control cells **(+DMSO)**. On the contrary, a limited, yet statistically significant drop of the F-actin density is detected in hCD4+ and hCD8+ T cells, but not in NK92 cells. **(c)** Live cell atomic force microscopy probing (AFM) shows rigidification of exponential phase hCD4+ T cells upon the build-up of microtubules during UR214-9-mediated septin inhibition. (**1**) - Measured T cell mechanical rigidities as the effective cell surface tension. Both UR214-9- and Taxol-induced microtubule stabilization result in the T cell rigidification. (**2**) - Schematic representation of the experimental settings and basic force configurations for live T cell mechanical probing. *Note that suppression of actomyosin contractility with Blebbistatin co-treatment during **+UR214-9** or **+Taxol** treatments does not soften T cell down to the baseline level **(+DMSO)**, indicating the T cell rigidification is a combined effect of the cortical contractility and passive mechanical resistance of the enhanced microtubule apparatus. Note also that blebbistatin co-treatments do not always result in the statistically significant T cell softening within all ±Blebbistatin co-treatment pairs (i.e. **±Blebbistatin**, **+UR214-9±Blebbistatin**, and **+Taxol±Blebbistatin**) due to the presence of the relatively large, mechanically resistant nuclei (∼60-70% of T cell volume), and dense microtubules (**+UR214-9** and **+Taxol** treatments). Thus, actomyosin contractility does not always significantly contribute to the modulation of T cell rigidity, yet microtubules do. Similarly, pharmacologically-induced disintegration of the F-actin network **(+Latrunculin A)** shows no statistically significant T cell softening due to the presence of the underlying nucleus*.

### Amoeboid-to-mesenchymal-like transition of T cells enhances microtubules

The unprecedented morphological conformity of T cells to the adhesion cues raises the question of mechanisms of observed mesenchymal-like transition. To answer the posed question, we note that the septin GTPase inhibition in T cells is characterized by a visibly increased microtubules’ density and length **(Figures 2a** and **3a**, **+DMSO** *vs.* **+UR214-9**; **Figure SI4a-1)**. To confirm and quantify the structural enhancement of microtubules, we estimate and compare microtubule density in control **(+DMSO),** and UR214-9-treated live hCD4+, hCD8+ T cells, and NK92 lymphocytes **(Figure 3b)** by flow cytometry using SiR-Tubulin, a non-toxic cell-permeable dye that only binds and stains the filamentous tubulin, *i.e.* - microtubules. The results show that hCD4+, hCD8+ T cells, and NK92 lymphocytes display ∼60%, 45%, and ∼900% increase in microtubule network density, respectively, following septins inhibition **(+UR214-9**, 1 hour) compared to the controls **(Figure 3b, +DMSO)**. The inhibition of septin GTPases also induces partial loss of the F-actin structural density in hCD4+ and hCD8+ T cells but not in the NK92 lymphocytes **(Figure 3b)**, as measured with SiR-Actin, a cell-permeable F-actin dye.

For orthogonal cross-examination of the effect of UR214-9-mediated inhibition of the septin GTPase activity in the enhancement of microtubules, we proceed to direct measurements of the microtubules-associated mechanical effects onto the T cell rigidity by live cell atomic force microscopy (AFM). Our previous experiments with the taxol-mediated (paclitaxel) stabilization of microtubules demonstrated a substantial increase in the passive rigidity of T cells (8). Similarly, the UR214-9-mediated inhibition of the septin GTPase activity resembles taxol-mediated effects with respect to the increase of T cell mechanical rigidity **(Figure 3c**, **+UR214-9** *vs.* **+Taxol)**. Notably, co-treatments of T cells with actomyosin contractility inhibitor (*i.e.*, blebbistatin) with either UR214-9 or taxol do not soften the lymphocytes to the baseline rigidity observed for a sole blebbistatin treatment **(Figure 3c**, **±Blebbistatin** *vs.* **+UR214-9±Blebbistatin** *vs.* **+Taxol±Blebbistatin)** or the control condition **(+DMSO)**. Thus, our results indicate that the observed mechanical rigidification of T cells upon inhibition of septin’s GTPase activity is indeed mediated by the enhanced density of the microtubules network that adds to the passive T cell mechanical rigidity.

### Inhibited septins stabilizes microtubules *via* MAP4- and HDAC6-mediated signaling

The detected stabilization of microtubules may rely on multiple synergized contours of signaling that interconnect septins, microtubules, and actomyosin components **(Figure 4a)**. For example, intact (*i.e.*, GTPase-active) septin-7 protein suppresses the microtubule-stabilizing protein (MAP) MAP4 (28) while maintaining the downstream HDAC6-mediated deacetylation of microtubules (49). Thus, the effects of septin-7 inhibition converge into the stabilization of the microtubule network *via* both the enhancement of microtubule acetylation **(Figure 4a-1**; **Figure SI4a-2)** and enhancement of MAP4-to-MT binding **(Figure 4a-1**; **Figure SI4b-1)** that causes MAP4-induced microtubule detyrosination in T cells (50) **(Figure 4a-1)**.

**Figure 4.**
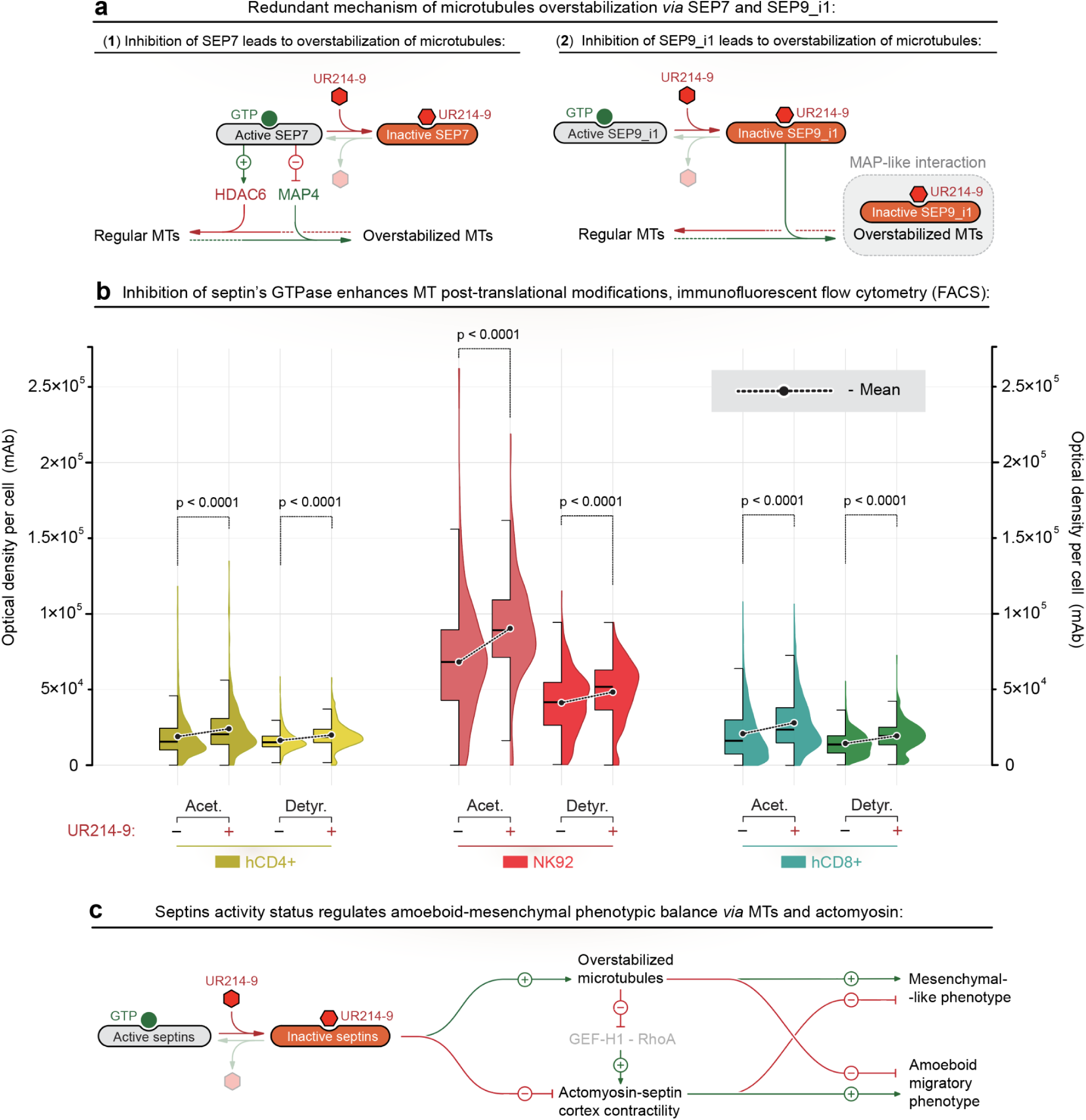
Septin-linked signaling pathways converge on microtubule stabilization during amoeboid-to-mesenchymal-like transitioning in lymphocytes. **(a)** Septin-7 (SEP7) and septin-9 (*i.e.*, septin-9, isoform-1, SEP9_i1) form parallel signaling circuits converging on the microtubule stabilization. **(1)** - GTPase activity of SEP7 supports the dynamic instability of microtubules both by maintaining microtubules deacetylation by HDAC6, and by deactivation of MT-stabilizing MAP4. Inhibition of SEP7 GTPase activity with the UR214-9 leads to the loss of HDAC6 activity and activation of MAP4, resulting in the stabilization of microtubules *via* enhanced post-translational acetylation of microtubules and MAP4 binding. **(2)** SEP9_i1 has a dual role as a septin and as an microtubule-associated protein *via* its MAP-like motif. Upon inhibition of its GTPase activity with UR214-9, released SEP9_i1 translocates onto the microtubules *via* its MAP-like motif, causing the SEP9-induced MAP-like microtubule-stabilization. Thus, UR214-9-treated cells can stabilize the microtubules by converging SEP7 and SEP9_i1 signaling circuits, suggesting that both septins form a redundant mechanism for regulation of microtubule dynamics. **(b)** Inhibition of septin’s GTPase activity with UR214-9 in hCD4+, hCD8+, and NK92 lymphocytes leads to the increase of post-translational microtubule modifications. Paired flow cytometry analysis of control and treated color-coded cells stained in mixed suspension shows both increase of tubulin acetylation **(acet.)** and detyrosination **(detyr.)** that indicate stabilization (*i.e.*, elongation and densification) of the microtubules (Figure 3b). **(c)** Septin GTPase activity regulates lymphocytes’ amoeboid-to-mesenchymal-like transition *via* microtubules and actomyosin. Stabilization of the microtubules *via* septin inhibition **(+UR214-9)** enhances lymphocytes’ mesenchymal phenotype, while it also suppresses the actomyosin contractility by sequestering the cytoplasmic GEF-H1 in the enhanced microtubule network. Suppression of the actomyosin contractility contributes towards the weakening of the amoeboid cortical dynamics and shifts cells towards the mesenchymal-like behavior.

Alternatively, SEPT9_i1 isoform directly binds to microtubules *via* its MAP-like motif (29, 30, 43), stabilizing the microtubules by preventing the MT filaments growth catastrophes (29), either indirectly or directly by intrinsic septin-9 MAP-like activity (30). Thus, UR214-9-mediated GTPase inactivation and release of septin-9 from its filamentous, actomyosin-associated form results with septin-9 translocates onto the microtubules *via* an intact MAP-like motif, stabilizing the microtubule apparatus **(Figure 4a-2**; **Figure SI4b-2)**. Indeed, GTPase activity is dispensable for septin-9’s MAP-like MT-binding and MT-stabilizing activities in MDA-MB-231 cells, where UR214-9 induces *en masse* septin-9 translocation from the stress-fibers to the MTs (37), accompanied by a visible enhancement of microtubule density in non-immune cells **(Figures 1c** and **SI2)**, as well in immune cells **(Figure SI4a-1)**. The suggested mechanism of septin-9’s MAP-like MT-binding and MT-stabilizing activities is supported by the effects of previously observed septin-9→kinesin signaling pathway that cause kinesin-driven microtubules curling, looping, and circularization into the microtubule rings **(Figure SI2**, **+UR214-9**, *arrows*), reported for various types of cells (37).

To explore the suggested septin-based signaling model, we proceed to the analysis of post-translational modifications of microtubules in response to the UR214-9-mediated septin’s GTPase inhibition. Flow cytometry analysis of post-translational modifications of microtubules in the control **(+DMSO)** and UR214-9-treated **(+UR214-9)** cells shows an unanimous ∼25% increase in both microtubule acetylation and detyrosination for hCD4+, hCD8+ T cells, and NK92 cells upon UR214-9-mediated inhibition of septin’s GTPAse activity **(Figure 4b)**. Similarly, the confocal microscopy of the spread hCD4+ T cells displays a substantial increase of the visible fraction of the acetylated microtubules, *i.e.*, from short MTs located only at the MTOC towards the long and dense MT filaments throughout the entire T cell volume **(Figure SI4a-2)**.

Our observations confirm the stabilization of microtubules, suggesting that inhibition of the septins may cause the amoeboid-to-mesenchymal transition in T cells *via* the following, synergized and complementary processes: (1) MT network enhancement (51) followed by (2) actomyosin contractility weakening (8) **(Figure 4c)**, attributed both to an increase of the microtubule network density that sequesters the cytoplasmic GEF-H1 (52), an upstream activator of RhoA-mediated pathway of actomyosin contractility (53), and to the loss of the septin-mediated stabilization of contractile actomyosin (38, 54).

### Mesenchymal-like T cell behavior depends on dynein-generated forces

Mechanistically, microtubules converge into a singular force transmission system with the F-actin cytoskeleton *via* force-generating dyneins (23, 24) and dynein cofactors, such as dynactin, that directly interlink dynein motors to the actomyosin network (24, 55–57). These dynein cofactors allow transmission of dynein-generated forces within microtubules to the F-actin-integrated adhesion receptors, which in turn facilitate forces for mechanosensing of the adhesion cues and guide the spreading of T cells during mesenchymal-like migration. Since dynein-MT affinity and processivity are enhanced by microtubules acetylation (58, 59) and detyrosination (60), we propose that septins inhibition increases the contribution of dynein-generated mechanical forces and stimulates T cell-to-ICAM1 adhesion, spreading, and migration. Moreover, if combined with the partial loss of the stable F-actin during septin GTPase activity inhibition **(Figure 3b**, **+DMSO** *vs.* **+UR214-9)**, the role of the enhanced microtubule network and dynein-generated forces is expected to increase upon UR214-9-induced amoeboid-mesenchymal transition **(Figure 5a)**. In this model enhanced elongated MTs are utilized as a system of long-distance dynein-generated forces transmission that effectively replaces actomyosin contractility and enables T cell adhesion and spreading along the structurally complex adhesion guidance cues (*i.e.*, ICAM1). We argue that upon septin inhibition the diminished 3D cortical actomyosin tension **(Figure 5a-1)** no longer interferes with the forces within the 2D T cell-adhesion interface. Combined with the dynein-generated tension that conforms along the microtubular cables network that is inherently congruent to the adhesion configuration **(Figure 5a-2)**, the induced effects switch from the amoeboid random walk towards the T cell towards the adhesion-based contact guidance.

**Figure 5.**
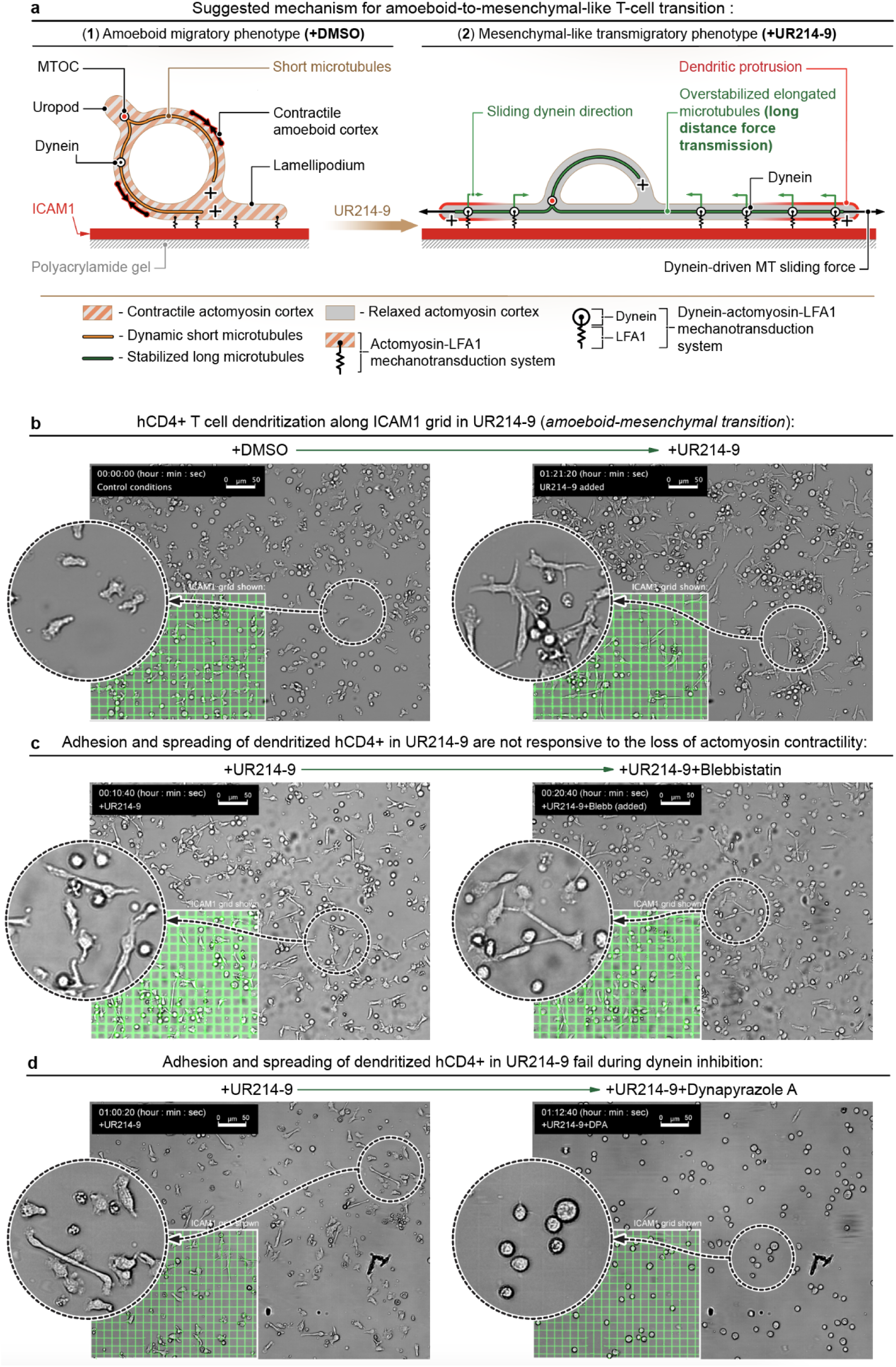
Lymphocytes that undergo amoeboid-to-mesenchymal-like phenotypic transitioning (hCD4+ cells, +UR214-9) resist the actomyosin inhibition, but are sensitive to inhibition of the dynein motors. **(a)** Schematic representation of the proposed mechanism for T cell transitioning from amoeboid **(+DMSO)** towards mesenchymal-like **(+UR214-9)** phenotype. (**1**) - Amoeboid T cells are largely unresponsive to the guidance cues used in the flat 2D systems because of complex shape-changing dynamics and highly contractile actomyosin cortex that dominate over the flat 2D adhesion ‘footprint’. (**2**) - Inhibition of septins activity suppresses cortical dynamics of actomyosin in amoeboid T cells. Simultaneously, inhibition of septin GTPase activity stabilizes microtubules that enhances processivity of dynein motors and transmission of dynein-generated forces along microtubule cables, which facilitates integrin-dependent mesenchymal-like T cell migration along the adhesive ICAM1 cues. **(b)** Amoeboid-to-mesenchymal-like transitioning of hCD4+ T cells. Lymphocytes form dendritic-like protrusions and demonstrate congruent alignment to the ICAM1 grids **(+DMSO→+UR214-9**, **Movies 4** and **5)**. **(c)** Mesenchymal-like hCD4+ T cells aligned to the ICAM1 grids **(+UR214-9)** are unresponsive to co-suppression of the actomyosin contractility by blebbistatin **(+UR214-9→+UR214-9+Blebb**, **Movie 6)**. **(d)** Mesenchymal-like hCD4+ T cells aligned to the ICAM1 grids **(+UR214-9)** are highly sensitive (cells collapse) to co-suppression of dynein motors with Dynapyrazole A **(+UR214-9→+UR214-9+DPA**, **Movie 7)**.

To examine the proposed mechanism of microtubule stabilization and upregulation of dynein-generated forces *via* MAP4-, HDAC6- and SEP9-mediated pathways **(Figure 4b)**, we present hCD4+ T cells to the ICAM1-based 2D contact guidance cues such as orthogonal grids **(Figure 3a)**. The hCD4+ T cells display a non-spreading amoeboid morphology and meandering ‘random walk’ migration behavior in control conditions **(Figure 5b**, **+DMSO, Movie 4)**, regardless of the geometry of the underlying ICAM1 grid. Septin inhibition with UR214-9 abruptly (*i.e.*, bi-phasically) switches T cells to their ‘dendritic’ mesenchymal-like spreading along the ICAM1 grid within the first 30 minutes of treatment **(Figure 5b**, **+DMSO**→**+UR214-9**, t=30 min, **Movie 5)**.

Subsequently, we question whether actomyosin or microtubules and dyneins ensure T cell mesenchymal-like adhesion and spreading along the ICAM1 grids in the presence of UR214-9. Intriguingly, suppression of the actomyosin contractility in the UR214-9-treated hCD4+ T cells **(+UR214-9**→**+Blebbistatin+UR214-9)** does not affect structural compliance and migrational guidance response of T cells to ICAM1 grids **(Figure 5c**, **+UR214-9**→**+UR214-9+Blebbistatin**, **Movie 6)**. Thus, our results indicate that actomyosin contractility is dispensable for mesenchymal-like migration and contact guidance of hCD4+ T cells upon inhibition of septins GTPase activity.

Next, we test our hypothesis that dynein activity within enhanced microtubules **(Figures 3a-c** and **4b)** may, at least partially, substitute actomyosin contractility in hCD4+ T cells subjected to the acute septin inhibition with UR214-9 **(Figure 5a-2)**. For that we target the dynein activity in the UR214-9-pretreated hCD4+ T cells, spread on the ICAM1 grids, by co-treatment with Dynapyrazole A (61) **(Figure 5d**, **+UR214-9**→**+UR214-9+Dynapyrazole A)**. Strikingly, unlike acute actomyosin contractility suppression, inhibition of dynein activity indeed abruptly truncates dendritic adhesion and spreading of UR214-9-pretreated hCD4+ T cells along the ICAM1 grids **(Movie 7)**. Thus, dynein activity appears indispensable for mesenchymal-like adhesion and spreading of T cells upon septin inhibition.

The subsequent washout experiment demonstrates the reversibility of the mesenchymal-like cytoskeleton dynamics, *i.e.,* a recovery of the T cell amoeboid migration **(Movie 8**, *washout*). Importantly, the separate hCD4+ T cell treatment with either sole blebbistatin or sole dynapyrazole A (*i.e.*, in the absence of UR214-9 septin inhibitor) does not show any T lymphocyte transitioning towards the mesenchymal phenotype **(Movie 9**, **+DMSO→+Blebbistatin**; **Movie 10**, **+DMSO→+Dynapyrazole A)**, indicating the key role of septin-centered signaling pathways in the regulation of microtubules and F-actin cytoskeleton components during T cell amoeboid-to-mesenchymal transition.

Furthermore, to challenge our conclusions on the importance of the microtubular apparatus enhancement for the amoeboid-to-mesenchymal transition of T cells, we aim to bypass septin inhibition and directly stabilize microtubules. For that purpose, we induce microtubule hyperacetylation and stabilization by directly suppressing HDAC6 with tubacin (62) **(+Tubacin,** 25 µM, 1 hour). Our results show that tubacin-treated T cells partially replicate the UR214-9-induced mesenchymal-like mode transition, *i.e.,* by phenocopying the dendritic-like spreading along the ICAM1 grids **(Figure SI5a**, **+DMSO** vs. **+Tubacin)**. The dendritic-like spreading of tubacin-treated T cells is accompanied by a significant enhancement of the microtubule acetylation that expands from the limited MTOC-associated acetylated tubulin cluster **(Figure SI4b**, **+DMSO**, *arrows*) to the dense and elongated acetylated microtubule network **(Figure SI5b**, **+Tubacin)**. Based on these observations, we conclude that partial amoeboid-to-mesenchymal transition of T cells can be achieved by microtubule hyperacetylation.

## DISCUSSION

Amoeboid T cells are typically non-responsive to the adhesive guidance cues used in the *in vitro* 2D systems (8, 12), despite the active adhesiveness to the ICAM1 *via* LFA1 integrin receptors (6, 12). The resulting meandering or ‘random’ walk migration behavior of T cells emphasizes that complex amoeboid shape-changing dynamics of the entire actomyosin cortex dominates over the reduced T cell’s 2D adhesion ‘footprint’ (6, 8). This actomyosin dynamics allows T cells to redistribute cortical stresses regardless of the configuration of the underlying 2D adhesive guidance cues (8, 51). However, the exact mechanisms that allow the adhesion-free cortical dynamics to dominate over the 2D adhesion are unclear (7). For example, shifting T cells away from the amoeboid phenotype with actomyosin contractility inhibitors does not enhance their contact guidance along the flat (2D) adhesion guidance cues, and poor 2D contact guidance remains largely indifferent to the modulation of microtubules→GEF-H1→RhoA signaling axis in T cells (8).

In this study, we experimentally demonstrate that acute pharmacological inhibition of septin’s GTPase activity induces the amoeboid-to-mesenchymal transition of T cells, characterized by distinct contact guidance on ICAM1 substrates. Based on our tests, we propose a model **(Figure 5a, 1 and 2)** describing how pharmacological inhibition of septins triggers the transition between (1) actomyosin-driven amoeboid migration **(Figure 5a-1)**, and (2) dynein-driven mesenchymal-like migration **(Figure 5a-2)**.

Specifically, our model suggests that the inactivation of septins must reduce the dynamics of the amoeboid actomyosin cell cortex to the level that allows for migration compliance to the 2D adhesive guidance cues. Indeed, septin inhibition with UR214-9 causes septin release from actomyosin and relaxation of the amoeboid cortex (63). Similarly, microtubule stabilization, *e.g.,* induced by septin inhibition, causes uptake of cytosolic GEF-H1 with a subsequent reduction of the RhoA→actomyosin contractility signaling axis (8, 51). Particularly, we show that inactivated septins may stabilize microtubules *via* MAP4 and septin-9 MAP-like pathways, hence facilitating the microtubule-mediated uptake of GEF-H1 from the cytoplasm and causing additional suppression of actomyosin contractility **(Figure 4c)**.

Additionally, our model proposes that the septins-mediated stabilization of microtubules enhances the processivity of dyneins within the microtubules and compensates for the diminished contribution of actomyosin contractility during the adhesion-based LFA1 activation, *i.e.,* in the absence of the stabilizing septin-actomyosin interactions (54, 63–65). For example, like other integrin receptors, ICAM1 receptor LFA1 is a mechanosensor (66–68). Therefore, dyneins in the UR214-9-treated T cells, during contact guidance on ICAM1 lines, must generate sufficient tension within LFA1 β2 subunit clusters to activate and sustain cell adhesion, spreading, and mesenchymal-like migration (69). Notably, for mesenchymal cells in a low actomyosin contractility state, stabilization and elongation of microtubules is critically important for developing and maintaining the dendritic protrusions along the collagen type-1 contact guidance cues (39, 46, 70, 71). Moreover, based on our observations, we conclude that partial amoeboid-to-mesenchymal transition of T cells can be achieved by microtubule hyperacetylation.

We note that the orthogonal AFM tests indicate that the observed mechanical rigidification of T cells upon inhibition of septin’s GTPase activity is mediated by the enhanced density of the microtubules network. While we have previously demonstrated that nocodazole-induced T cell rigidification was entirely reversed by the following Blebbistatin co-treatment (8), co-treatments of T cells with actomyosin contractility inhibitor, *i.e.*, Blebbistatin, with either UR214-9 or Taxol does not soften the lymphocytes to the baseline rigidity observed for a sole Blebbistatin treatment. Therefore, AFM measurements indicate that enhanced microtubules compensate for the diminished mechanical contribution of actomyosin to the cell rigidity.

Our previous report showed that dynein activity in mesenchymal cells regulate the spreading of dendritic-like cell protrusions and topography-mediated contact guidance (37). We also demonstrated that dyneins actively pull the microtubules into the adhesion sites to facilitate cell-substrate steric interactions and cell alignment (46), which is similar to dynein’s role in fibroblast’s migration upon suppression of the RhoA-dependent actomyosin contractility (72). These reports support our data and conclusion that inhibition of septins GTPase activity triggers amoeboid-to-mesenchymal-like transition and contact guidance in T cells *via* upregulation of dynein-driven forces within the enhanced microtubules. These dynein-driven forces produce compensatory tension substantial for activating LFA1 withoutsufficient actomyosin tension, which becomes diminished or even dispensable during mesenchymal-like migration.

In summary, we demonstrate that septin’s GTPase activity provides the On-Off switch of integrin-dependent guided migration of T lymphocytes. Acute inhibition of septins GTPase activity activates propulsion of T cell by microtubule-and dynein-based force-producing systems during mesenchymal-like migration along the external adhesion cues. These results suggest that T lymphocytes rely on septins, adapting for better migration toward sites of tissue inflammation and potential treatment sites in therapy settings.

## MATERIALS AND METHODS

### Key resources table

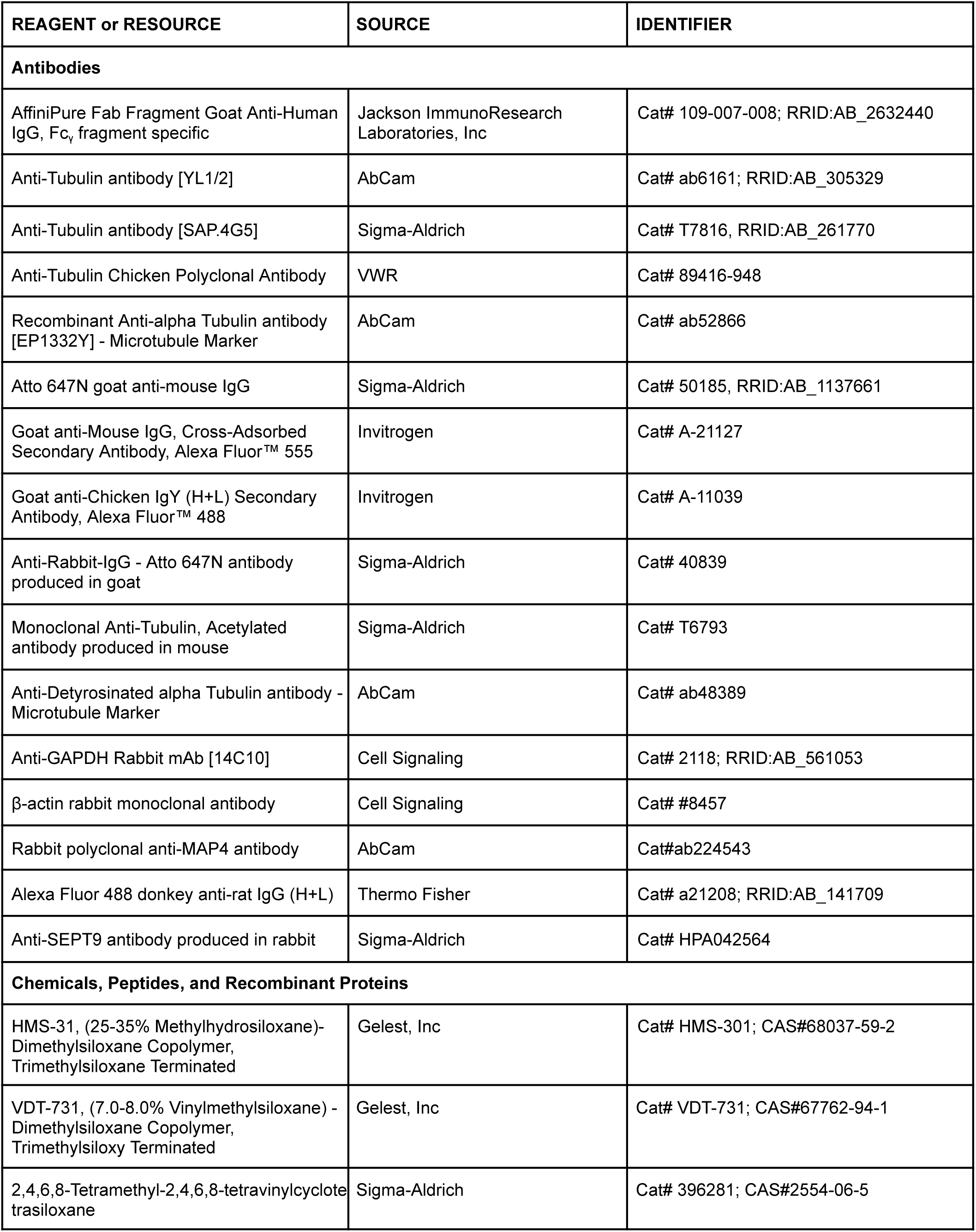
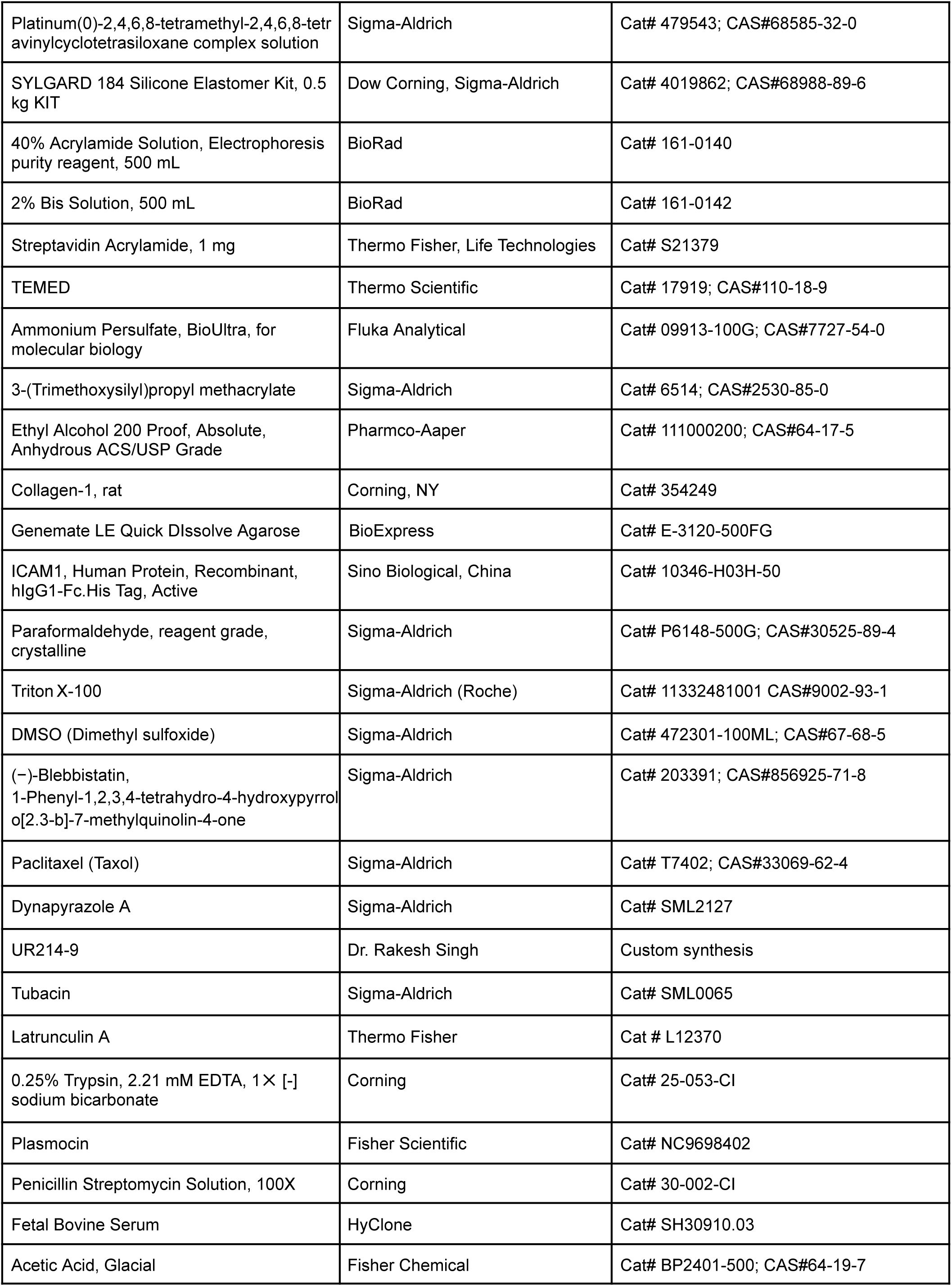
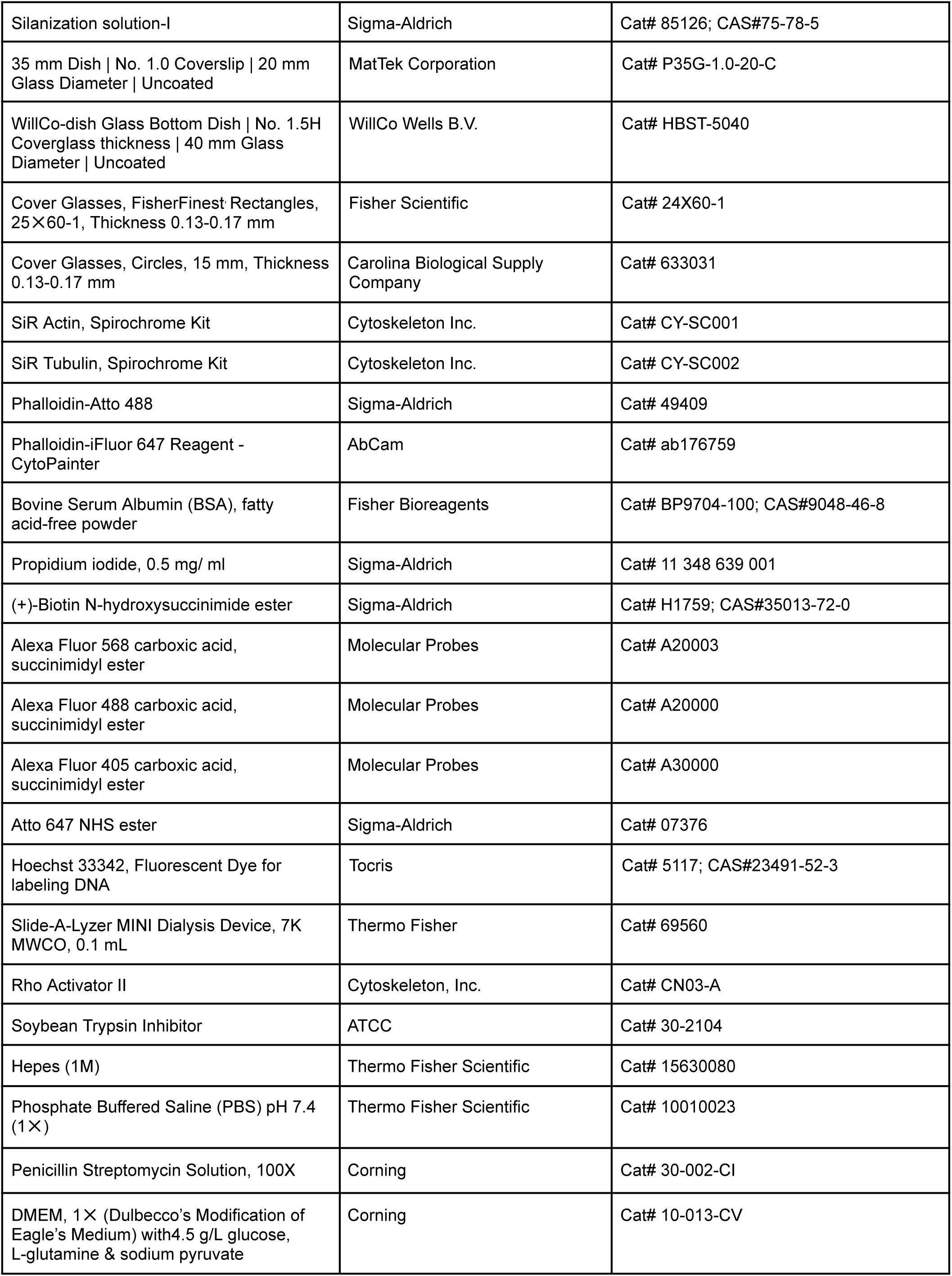
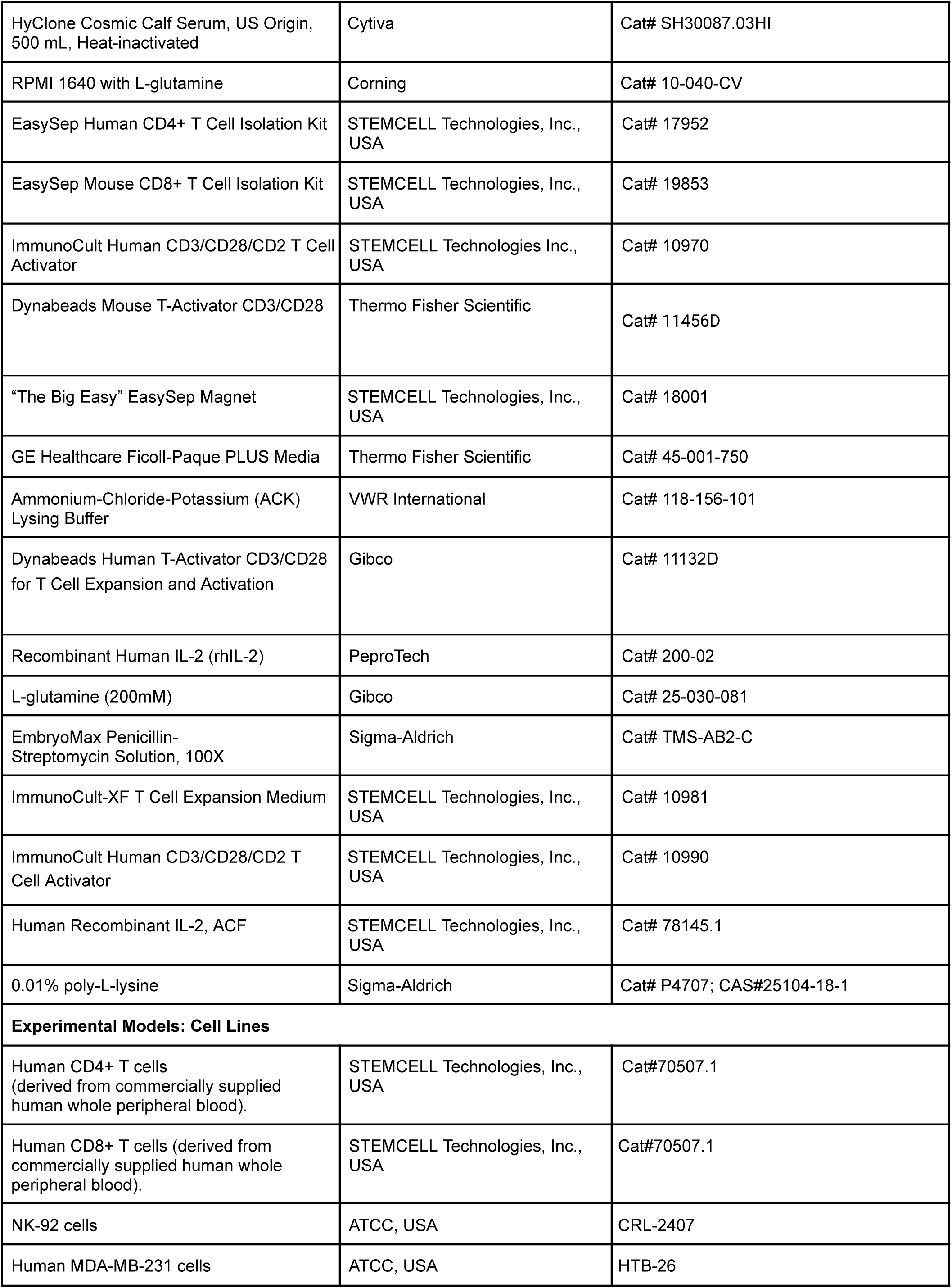
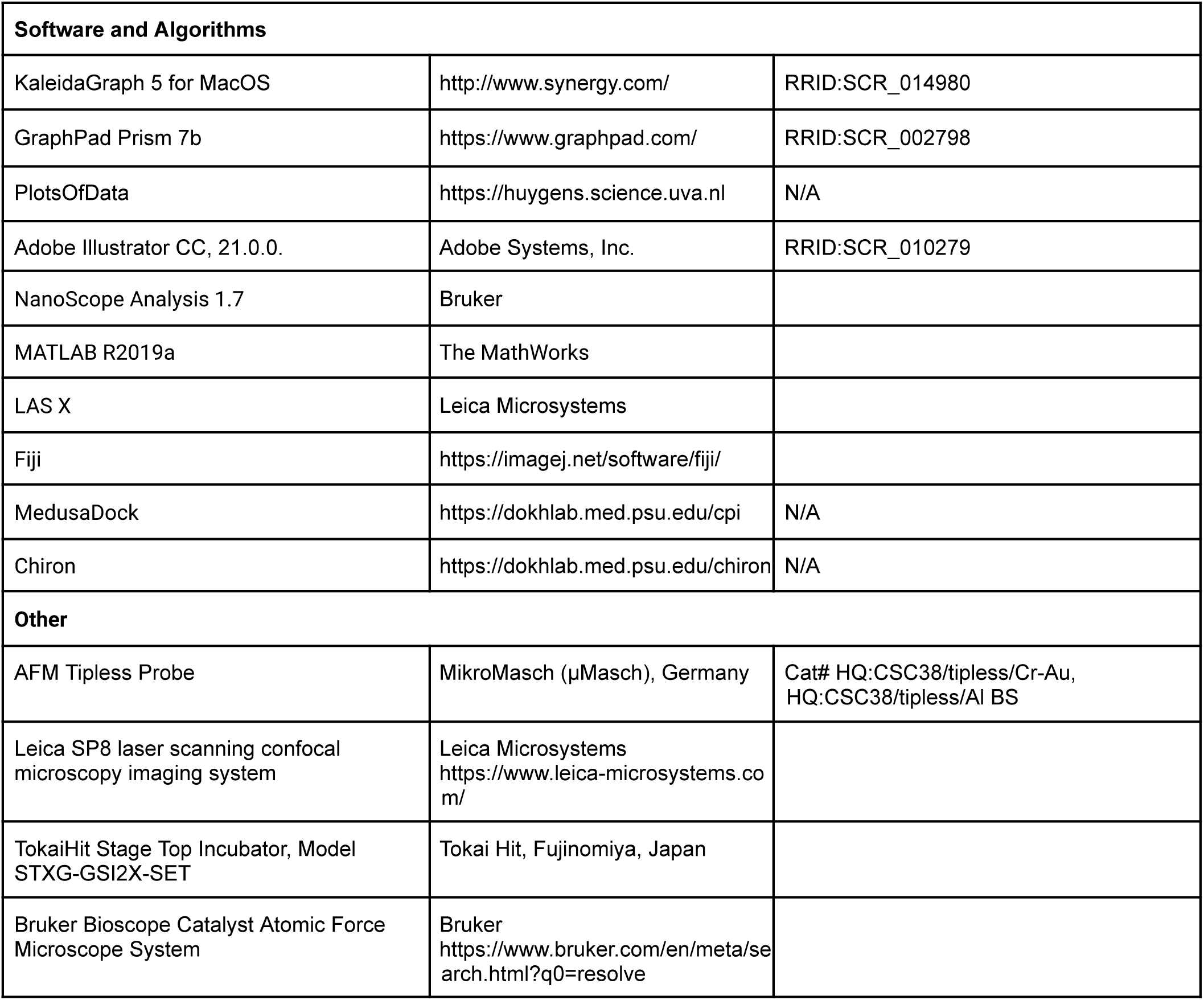

### Cell experiments

We maintained human MDA-MB-231 cells (ATCC® HTB-26™) in DMEM with 4.5 g/L D-glucose, L-glutamine,110 mg/L sodium pyruvate (Corning Cellgro®, Cat# 10013CV) and 10% heat-inactivated FBS (HyClone®, Cat# SH30071.03H) at 37°C in 5% CO2. For NK-92 cells (ATCC® CRL-2407), we used ImmunoCult-XF T Cell Expansion Medium (STEMCELL Technologies Inc., USA) with the addition of 1 μg/L of human interleukin 2 (IL-2, STEMCELL Technologies Inc., USA).

Primary human CD4+ and CD8+ T cells were isolated from commercially available whole human blood (STEMCELL Technologies Inc., USA, Cat# 70507.1) with EasySep Human CD4+ or CD8+ T Cell Isolation Kits (STEMCELL Technologies Inc., USA). Unless otherwise indicated, all immune cells were cultured, activated and expanded in ImmunoCult-XF T Cell Expansion Medium (STEMCELL Technologies Inc., USA) with the addition of ImmunoCult Human CD3/CD28/CD2 T Cell Activator and Human Recombinant Interleukin 2 (IL-2, STEMCELL Technologies Inc., USA) as per STEMCELL Technologies Inc. commercial protocol, at 37°C in 5% CO2. T cells were expanded only to the exponential growth phase to avoid cell exhaustion and/or quiescence.

### High precision micropatterning

An instructive and detailed protocol for indirect, high-definition, submicron-scale polyacrylamide gels micropatterning with various proteins is described in the original report (73). Briefly, collagen type-1 micropatterning at the micron-and submicron scale spatial precision is technically complicated by the capillary and van-der-waals effects that routinely induce a collapse of the microstamp bas-reliefs, made of conventional soft regular PDMS (rPDMS, polydimethylsiloxane), onto the printed glass surface. In order to preclude the microstamps from undesirable collapsing we replaced rPDMS stamping bas-relief with a 0.5-0.8 mm-thick veneering microprinting surface, *i.e.* - hard PDMS (hPDMS) (46, 74), mounted on the 5-8 mm-thick cushioning rPDMS blocks. To cast the micro-printing surfaces from the commercially fabricated and passivated molding matrix of desirable design (Minnesota Nano Center, University of Minnesota), we coated molds with 0.5-0.8 mm-thick layer of hPDMS either manually with a soft paraffin spatula or on a spin coating device. The submillimeter hPDMS layer on the silicone molding matrix was then cured in a heat chamber (70°C, 30 minutes). Next, an ∼8 mm-thick layer of bubble-free rPDMS was cured atop of the hPDMS layer (rPDMS; 1:5 curing agent/base ratio, Sylgard-184, Dow Corning, 70°C for ∼1 hour). The cured micro-stamping composite was then peeled off the mold and cut into 1✕1 cm pieces for the further use.

In order to assemble the elastic ICAM1 or collagen type-1 micro-patterns, we first microprinted α-Fc FAB antibody fragment (Jackson Immunoresearch, USA, Cat# 109-007-008, RRID:AB_2632440) or α-collagen-1 rabbit pAb (AbCam, Cambridge, UK, Cat# ab34710; RRID:AB_731684) on clean “intermediate” glass surface, stripped from organic impurities by the prolonged thermal treatment (FisherFinest™ Premium Cover Glass; #1, Cat# 12-548-5 P; 450°C, baked 20 hours in the furnace). For that, the cognate antibody for collagen-1 or an antibody fragment for ICAM-1-hFc were covalently conjugated with biotin, ((+)-biotin *N*-hydroxysuccinimide ester, Sigma-Aldrich, Cat# H1759; as per the manufacturer’s protocol) and a fluorescent Alexa Fluor tag (Alexa Fluor® succinimidyl esters, Invitrogen™, Molecular Probes®, Cat# A20000, Cat# A20003; as per commercial protocol). For that, the micro-stamps were coated with the 0.2 mg/mL α-collagen-1 antibody (or anti-Human Fc FAB) PBS solution for 40 min at 37°C in a dark humid chamber, then gently rinsed in deionized water, and dried under a jet of argon or nitrogen immediately prior to the micro-contact soft lithography procedure. Second, glass-bottom 35 mm cell culture dishes (MatTek Corp., Ashland, MA, Cat# P35G-1.0-20-C) were activated with 3-(trimethoxysilyl)propyl methacrylate (Sigma-Aldrich, Cat# 6514, as per commercial protocol) to ensure the covalent crosslinking between the glass surface and PAA gels. Third, 5 μL of PAA premix (polyacrylamide, see the “PAA elastic gel premix” section) of the projected rigidity G’ = 50 kPa(75) with the addition of the 5% streptavidin-acrylamide (ThermoFisher, Cat# S21379) were, avoiding any air bubbles formation, carefully ‘sandwiched’ between the activated glass surface and the micropatterned intermediate glass surface promptly after adding a curing catalyst (aminopropyltriethoxysilane, APS). The resulting ‘sandwiches’ with cured PAA were then incubated in deionized water (1 h, 20°C) for hypotonic PAA swelling that ensures gentle coverglass release from PAA gel. The resulting dish-bound PAA gels were checked for quality of the transferred fluorescent micropatterns on an epifluorescent microscope. Lastly, the anti-Human Fc FAB patterns on PAA were incubated with 10 µg/mL ICAM1-hFc solution (Sino Biological, Cat# 10346-H03H-50) and α-collagen-1 Ab micropatterns on PAA gels were incubated with 1 mg/mL rat monomeric collagen-1 (Corning, NY, Cat# 354249) in cold PBS (4 °C, 12 hours), then rinsed three times with cold PBS, and utilized for experiments.

### hPDMS formulation

To prepare the hPDMS mixture we used 3.4 g of VDT-731 (Gelest, Inc., Cat# VDT-731), 18 μL of Pt catalyst (Platinum(0)-2,4,6,8-tetramethyl-2,4,6,8-tetravinylcyclotetrasiloxane complex solution) (Sigma-Aldrich, Cat# 479543), and one drop of cross-linking modulator 2,4,6,8-Tetramethyl-2,4,6,8 -tetravinylcyclotetrasiloxane (Sigma-Aldrich, Cat# 396281). All components were thoroughly mixed in a 50 mL conical bottom centrifuge tube on the vortex mixer for at least 30 sec. Before use, we added 1 g of HMS-301 (Gelest, Inc., Cat# HMS-301) to the mixture, promptly mixed it for 30 sec on a vortex mixer, degassed in a high-speed centrifuge (3 minutes, 3000 rpm), and immediately used it for the mold coating. The detailed protocol is described elsewhere (74, 76).

### PAA elastic gel premix

For PAA premixes, we utilized 40% acrylamide (40% AA) base (BioRad, Cat# 161–0140) and 2% bis-AA (BioRad, Cat# 161–0142) cross-linker as described elsewhere.(77, 78) Additionally, streptavidin-acrylamide (Thermo Fisher, Cat# S21379) was added to the final concentration of 0.133 mg/mL to enable PAA gels cross-linking with biotinylated proteins of interest for the micropatterns transfer from the intermediate glass surface to the cured PAA gels. Briefly, for preparation of 50 µL of G’ = 50 kPa PAA gel premix, the components were mixed as follows: 40% AA - 15 µL; 2% bis-AA - 14.40 µL; 2 mg/mL streptavidin-acrylamide - 3.33 µL; 10X PBS - 5 µL; deionized milli-Q water - 11.17 µL; TEMED - 0.1 µL. The premix solutions were degassed and stored at 4°C before use. To initiate curing, 1 µL of 10% APS was added to 50 µL of PAA premix immediately before the PAA gel casting procedure.

### Synthesis of UR214-9

Equimolar mixture of 3-amino-2-fluorobenzotrifluoride (Combi-Blocks, Cat#QA-4188) and 2,6-dichloro-4-isocyanatopyridine (Toronto Research Chemicals, cat#159178-03-7) were stirred and heated in anhydrous Toluene at 85°C overnight. The separated UR214-9 was filtered and dried under vacuum in a dessicator. A portion of UR214-9 was detected in the Toluene layer, which was concentrated and purified by thin-layer chromatography using ethyl acetate and hexane as eluents. The pure product band was scrapped off the glass plate and UR214-9 was stripped from the silica gel using DCM+MeOH through a sintered funnel. The solvent was evaporated using a Buchi rotary evaporator to obtain UR214-9 as an off-white powder. The structure was confirmed by X-ray crystallography.

### Confocal microscopy

Cells were fixed with ice-cold (4°C) DMEM containing 4% paraformaldehyde (Sigma-Aldrich, Cat# P6148) for 20 minutes. PFA-fixed samples were rinsed in 1% BSA in PBS two or three times (5 minutes for each cycle), followed by 30 minute-long blocking-permeabilization treatment in 0.1% Triton X-100 (Sigma-Aldrich, Cat# T8787) solution with 10% BSA (Fisher, Cat# BP9704) in PBS (Thermo Fisher, Cat# 10010023). For immunofluorescence sample staining, we utilized primary antibodies diluted in 1% BSA PBS (all catalog numbers of antibodies are listed in Key Resources Table). The duration of the incubation with any of the listed primary antibody solutions was 1 hour at room temperature. Similarly, labelings with Alexa-Fluor-conjugated secondary antibodies (Thermo Fisher) were performed at their final concentration of 5 µg/mL for the duration of 1 hour in 1% BSA PBS at room temperature. After washing out the excess secondary antibodies, F-actin was stained either with fluorescent phalloidin or SIR-Actin for the duration of 1 hour. Chromatin was labeled with 1:1000 Hoechst solution (Tocris, Cat# 5117), added to the 90% Glycerol (Sigma, Cat# G5516) in 1x PBS mounting media.

High-resolution 3D and 2D imaging for cell morphometric analysis were performed on a Leica TCS SP8 laser scanning confocal microscope with LIAchroic Lightning system and LAS X Lightning Expert super-resolution capacity, 405, 488, 552 and 638 nm excitation diode lasers, with 40✕/1.3 oil immersion objective (Leica, Germany). The scanning settings were optimized with Nyquist LAS X function with HyD2.0-SMD excitation sensors, at the regular pinhole size of 0.85 AU, and the scanning frequency at 100 Hz. Each frequency channel was scanned in sequence in order to avoid the signal bleeding between the channels. Additionally, live cell imaging microscopy experiments were performed on epifluorescent Leica DMi8 microscope (Leica, Germany) equipped with a temperature (37°C), CO_2_ (5%), and humidity-controlled chamber at 20✕, 40X and 63X magnifications. Brightfield and fluorescence time-lapse images were obtained using LAS X software (Leica, Germany).

All analyses were performed automatically and/or manually utilizing Leica LAS X software (Leica, Germany) and the ImageJ/FIJI. Figures were composed using unmodified LAS X -generated TIFF images with Adobe Illustrator CC 2021 (Adobe).

### Flow cytometry

For analysis of microtubule and F-actin density in live cells, we stained CD4+, CD8+, and NK92 cells, control (DMSO) and UR214-9-treated (25 µM), with either SiR-Tubulin or SiR-Actin kit, respectively, following the commercial protocol, at 37°C in 5% CO2. Briefly, for SiR-Actin or SiR-Tubulin experiment, a cell aliquot was separated into eight tubes as follows: unstained control and UR214-treated cells, control and UR214-9-treated cells stained with propidium iodide (PI, 0.5mg/mL, 1:200, for 10 minutes prior to flow cytometry), control and UR214-9-treated cells stained with the SiR-reagent, control and UR214-9 cells stained both with PI and SiR-reagent. All UR-214-9-treated cells were incubated with UR214-9 for 1 hour before the addition of SiR-reagent, and for an additional hour after the addition of SiR-reagent. After the second hour of incubation, we added PI, where necessary, and proceeded with the flow cytometry analysis.

For analysis of post-translational modifications of microtubules in fixed cells, we stained mixed pre-labeled control (DMSO, Alexa405-negative) and UR214-9 treated (25 µM, Alexa405-positive) CD4+, CD8+, or NK92 cells with antibodies against acetylated (Monoclonal Anti-Tubulin, Acetylated antibody produced in mouse, Sigma-Aldrich, Cat# T6793) and detyrosinated (Anti-Detyrosinated alpha Tubulin antibody - Microtubule Marker, AbCam, Cat# ab48389) forms of tubulin. Briefly, each CD4+, CD8+, or NK92 cell aliquot was separated into two tubes - control and UR214-9-treated cells. After 2 hours of incubation with either DMSO or UR214-9, at 37°C in 5% CO2, we fixed control and UR214-9-treated cells with 4% PFA in 1x PBS for 20 minutes. After centrifugation at 300x g for 10 minutes, we removed excess PFA and additionally washed cells with 1 mL of 1x PBS. Fixed UR214-9 cells were resuspended in 0.1 mL of 1x PBS with 1:100 of 1 mg/mL Alexa Fluor 405 succinimidyl ester (Molecular Probes, Cat# A30000), and incubated for 1 hour at room temperature. Excess of unreacted ester was neutralized with equal volume of 10% BSA in 1x PBS for 30 minutes. Alexa405-labeled cells were washed with 1 ml of 1x PBS and mixed with the corresponding Alexa405-negative control cells for subsequent “one tube” permeabilization, blocking and staining with primary and secondary antibodies, i.e., goat anti-Rabbit Atto647N and goat anti-Mouse Alexa555 secondary antibodies, following the procedure described before for confocal microscopy staining. All washing steps for suspended cells were performed by centrifugation at 300x g for 10 minutes.

### Atomic Force Microscopy

Depending on the pharmacological drug treatment, expansion phase immune cells were incubated in Immunocult-XF culture media with the respective drug concentration (Blebbistatin: 50 µM, UR214-9: 50 µM, Taxol: 1 µM, Latrunculin A: 100 nM) for the given treatment condition for total times between ∼15 min to 75 min. The cells were incubated in Eppendorf tubes with their respective drug concentrations, followed by plating on a glass-bottom dish (WillCo Wells) precoated with 0.01% poly-L-lysine (Sigma-Aldrich). Plating was done ∼15 minutes before imaging to allow cells to weakly adhere to the glass-bottom dish surface and the drug to take maximum effect. AFM force spectroscopy experiments were performed using a Bruker BioScope Catalyst AFM system (Bruker) mounted on an inverted Axiovert 200M microscope (Zeiss) equipped with a confocal laser scanning microscope 510 Meta (Zeiss) and a 40x objective lens (0.95 NA, Plan-Apochromat, Zeiss). The combined microscope instrument was placed on an acoustic isolation table (Kinetic Systems). During AFM experiments, cells were maintained at physiologically relevant temperature 37°C using a heated stage (Bruker). A soft silicon nitride tipless AFM probe (HQ:CSC38/tipless/Cr-Au, HQ:CSC38/tipless/Cr-AlBS, MikroMasch) was used for gentle compression of spherical weakly-adherent cells. The AFM microcantilevers were pre-calibrated using the standard thermal noise fluctuations method, with estimated spring constants for microcantilevers used between 0.04–0.12 N/m. Immediately after probe pre-calibration, the AFM tipless probe was moved on top of a spherical cell. Five to ten successive force curves were performed on each cell. The deflection setpoint was set between 4-12 nm yielding applied forces between 0.16 to 1.44 nN.

All AFM force-distance curves measurements were analyzed using a custom-written MATLAB (The MathWorks) code to calculate the cellular surface cortical tension. For curve fitting, indentation depths between 0-600 nm were relatively consistent in yielding good fits (R2>0.75). Curves with poor fits R2<0.75 were discarded from the analysis. Additionally, we discarded noisy force curves and/or curves that presented jumps possibly due to cantilever and plasma membrane adhesion, slippage, or very weakly adhered moving cells.

Spherical cell cellular surface cortical tension (*T*; pN/µm) was calculated by fitting each recorded force-distance curve using the cortical tension method previously described that defines the force balance relating the applied cantilever force with the pressure excess inside the rounded cells and the corresponding actin cortex tension; 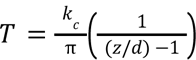, where *T* is the cellular surface tension, *k_c_* is the AFM cantilever spring constant, *z* is the Z-piezo extension, and d is the cantilever mean deflection.(79)

### *In silico* UR214-9-Septin docking energy simulation

The Swiss-Model (80), a homology modeling based protein structure prediction tool, was used to model the structures of Septin 2, 6, 7, and 9. Since these septins have crystal structures, the Swiss-Model was mainly used to complete the missing residues and atoms. Particularly, Septin 2, 6, 7, and 9 were modeled by using 6UPA, 6UPR, 6N0B, and 5CYO as the templates for homology modeling. The dimer structure of each of the septins is built by aligning two monomer structures to the corresponding positions in 2QAG, which is a septin oligomer structure. Next, the dimer structures were optimized by using Chiron (81). Two pockets were identified according to the positions of the ligands in 2QAG. The two pockets are at the dimerization interface of the dimer structures. MedusaDock (82, 83) was used to dock GTP, GDP, UR214-9, and FCF to the dimer structures of the septins. Docking poses were evaluated by MedusaScore, (84) a scoring function based on a custom physics-based force field. 1000 docking attempts were performed for each of the protein-ligand pairs, and the resulting docking poses were clustered by MedusaDock to select the centroid as the final candidate.

### Statistical analysis

Only pairwise comparisons as one-sided *t* tests between a control group and all other conditions are utilized to analyze the data, as well as between paired -Blebbistatin and +Blebbistatin co-treatment groups. Statistical analysis is performed using either KaleidaGraph 5 for MacOS (Synergy Software) or Prism 7b (GraphPad Software, Inc). The exact p values are indicated on the plots, unless the p<0.0001, *i.e.* below the cut-off lower limit for Kaleidagraph and Prism software. Sample size n for each comparison is reported in the corresponding plots (*i.e.* n reflects the number of measured individual cells). Number of replicates is 3, unless specified otherwise. Data are shown as box and whiskers diagrams: first quartile, median, third quartile, and 95% percent confidence interval.

## DATA AVAILABILITY STATEMENT

The authors declare that the data supporting the findings of this study are available within the paper, the supplementary information, and any data can be made further available upon reasonable request.

## ACKNOWLEDGEMENTS

E.D.T. and this work were supported by the Department of Pharmacology, Penn State College of Medicine *via* the startup funds. N.V.D. is supported by the NIH grant R35GM134864 and the Passan Foundation. D.T. is supported by the National Science Foundation (CMMI 1942561) and the National Institutes of Health (R01GM136892). A.S.Z., X.M., and A.M. were supported by the FDA Intramural Research Program of the Center for Biologics Evaluation and Research. A.X.C.R. and C.S. were supported by the National Institutes of Health (NIH) Intramural Research Program in the National Institute of Biomedical Imaging and Bioengineering (NIH grant # ZIA EB000094) and by the NIH Distinguished Scholars Program. We thank Christian Combs and Daniela Malide for the Light Microscopy Core support at the National Heart, Lung, and Blood Institute, NIH.

## MOVIE LEGENDS

**Movie 1.** Wide-field view of hCD4+ T cell migration on 2D (flat) ICAM1 parallel lanes micropattern during their transitioning from predominantly amoeboid ’random walk’ towards a mesenchymal-like contact guidance mode after the addition of UR214-9 septin inhibitor.

**Movie 2.** Detailed zoom-in view of hCD4+ T cell mesenchymal-like contact guidance along ICAM1 micro-lanes.

**Movie 3.** Wide-field view of the hCD4+ T cells’ contact guidance along ICAM1 micro-lanes during prolonged **+UR214-9** treatment (t > 1 hour).

**Movie 4.** hCD4+ T cells predominantly amoeboid ’random walk’ meandering on the flat 2D ICAM1 grids.

**Movie 5.** UR214-9-induced hCD4+ T cells mesenchymal-like spreading and formation of dendritic-like cell protrusions along the ICAM1 grid.

**Movie 6.** Non-responsiveness of UR214-9-induced mesenchymal-like hCD4+ T cell dendritic adhesion, spreading, and contact guidance dynamics to Blebbistatin-mediated suppression of actomyosin contractility.

**Movie 7.** Dynein contractility inhibition with Dynapyrazole A abrogates UR214-9-induced hCD4+ T cells mesenchymal-like adhesion, spreading, and migration along ICAM1 grids. *Note that the tested T cells are co-treated with UR214-9 and Blebbistatin to demonstrate dispensable status of actomyosin contractility during mesenchymal-like T cell contact guidance mode under UR214-9 treatment*.

**Movie 8.** Abrogation of hCD4+ T cell’s UR214-9-induced mesenchymal-like adhesion, spreading, and migration with Dynapyrazole A-induced dynein inhibition (no Blebbistatin co-treatment). Following washout demonstrates partial reversibility of treatment effects.

**Movie 9.** Inhibition of the actomyosin contractility **(+Blebbistatin)**, unlike UR214-9 treatment, does not induce mesenchymal-like ‘dendritic’ cell spreading.

**Movie 10.** Sole inhibition of the dynein contractility **(+Dynapyrazole A)** does not change hCD4+ T cell predominantly amoeboid migratory behavior on ICAM1 grids.

## SUPPLEMENTARY FIGURES

**Supplemental Figure SI1.**
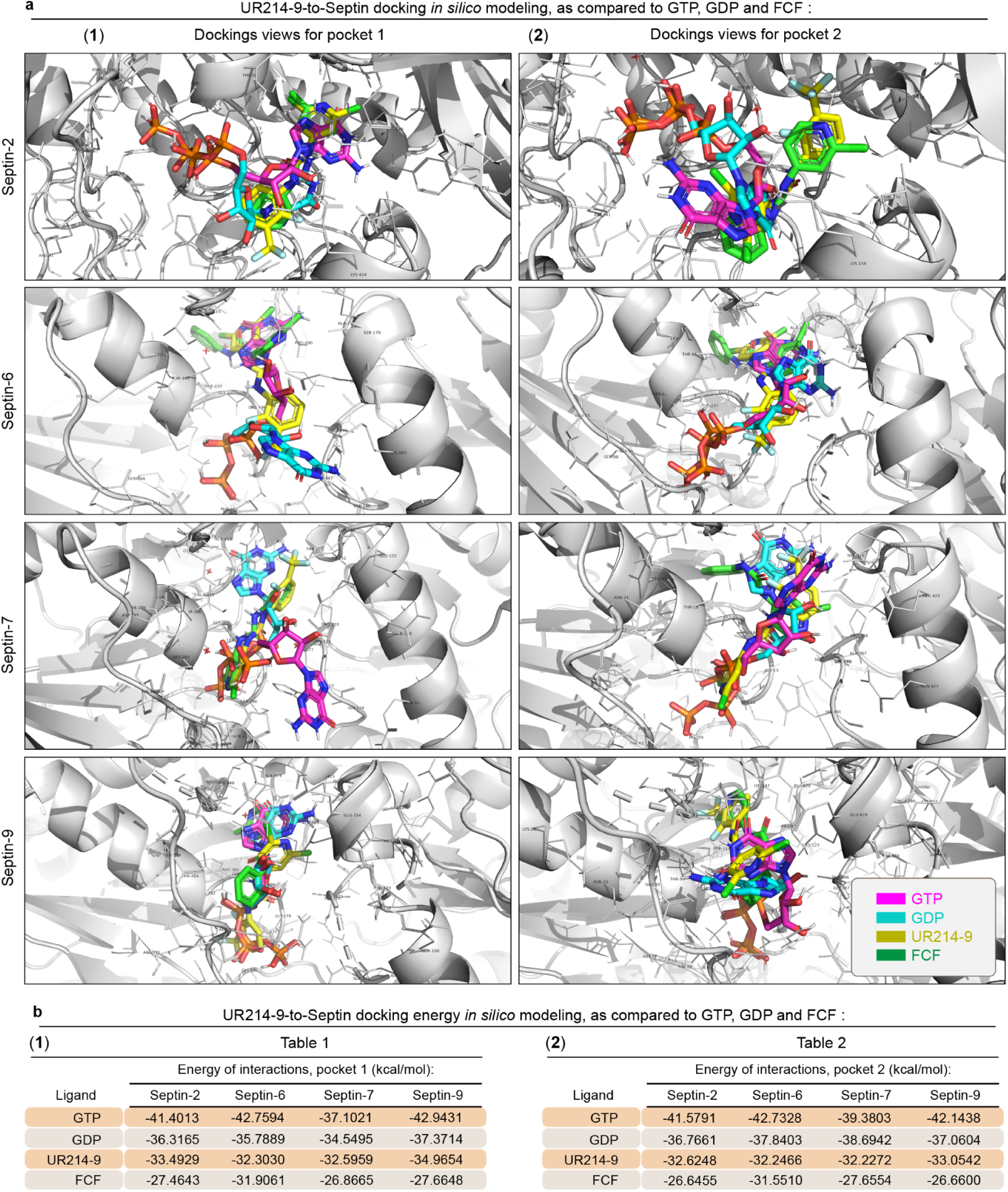
UR214-9-septin *in silico* docking simulations display competitive energies of interactions as compared to FCF, GTP and GDP. **(a)** *In silico* computational model for UR214-9 docking to septin-2, -6, -7 and -9’s GTPase site at the respective pockets 1 and 2 in septin molecules, as compared to FCF inhibitor and GTP. **(b)** *In silico*-calculated docking energies for GTP, GDP, UR214-9 and FCF.

**Supplemental Figure SI2.**
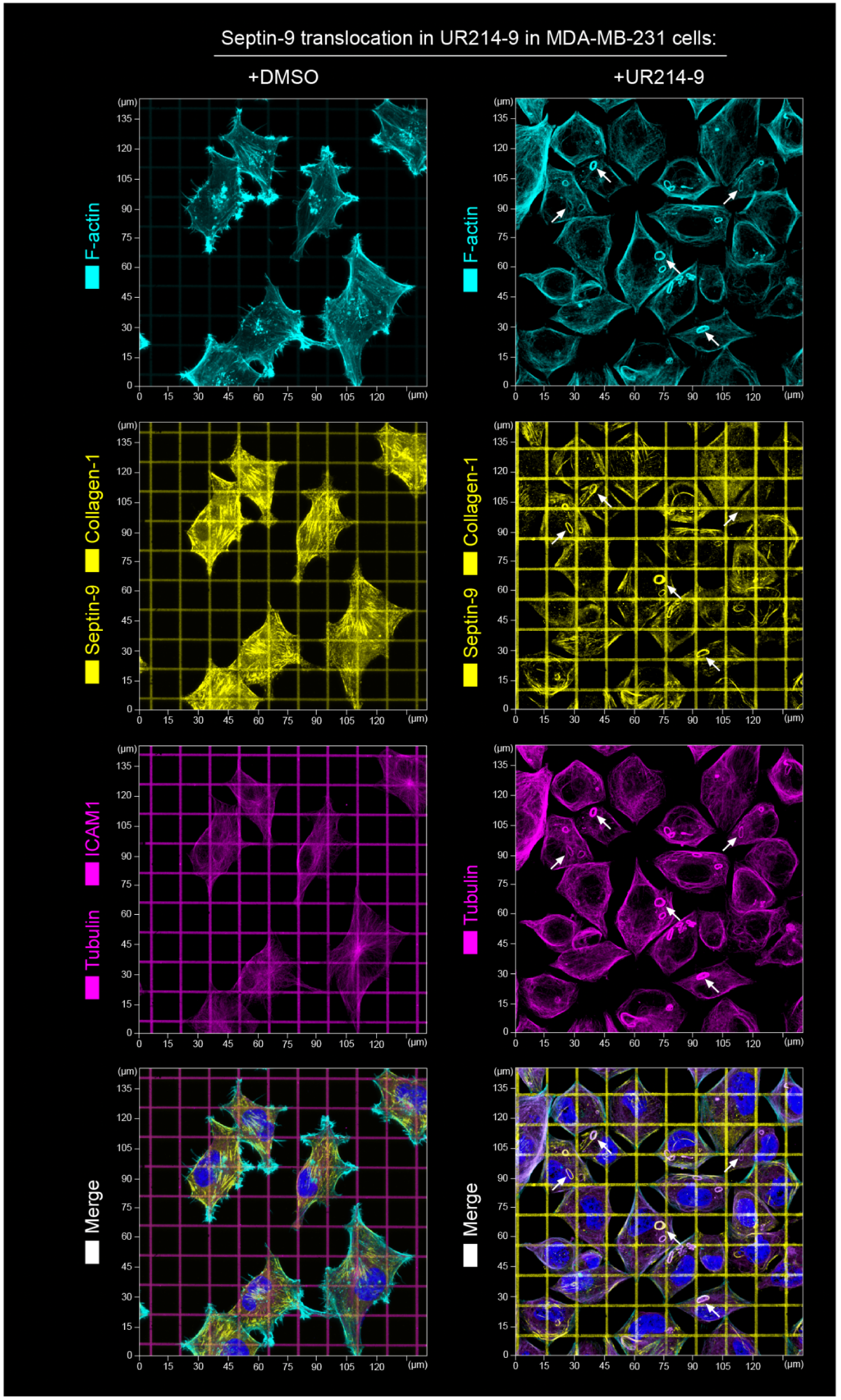
Septins inhibition (+UR214-9) in MDA-MB-231 cells on collagen (type I) grids results in translocation of septin-9 from the stress-fibers onto microtubules. *Left* - Septin-9 in vehicle-treated cells (+DMSO) colocalizes with linear F-actin structures (stress-fibers). *Right* - Septin-9 translocates onto the microtubules upon suppression of septins GTPase activity (+UR214-9, t≥1 hour). *Note that inactivation of GTPase activity if septins leads to the increased density of the MT network*. *Note that GTPase-inactive septin-9 acts as a microtubule-associated protein (MAP), stabilizing microtubules and inducing MT curling into the rings (arrows) (37)*.

**Supplemental Figure SI3.**
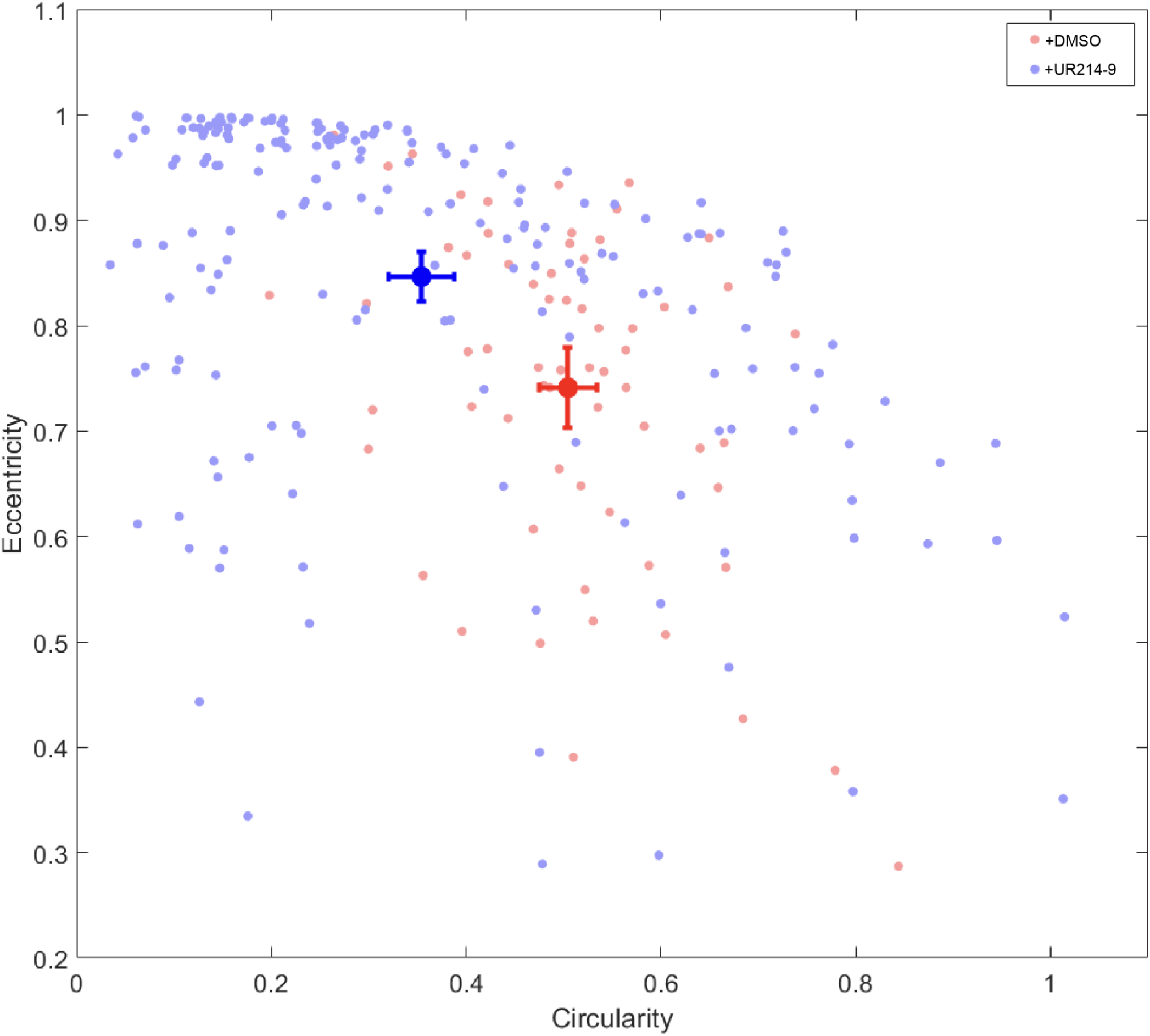
Analysis of hCD4+ T cells morphological response to the inhibition of the septins with UR214-9. A cross-correlated substantial eccentricity (T cell elongation) and circularity (4πS/L², where S is the projected cell area, L is projected cell perimeter) changes. *Note the increase in cell elongation (eccentricity) and decrease of T cell circularity upon treatment with UR214-9*.

**Supplemental Figure SI4.**
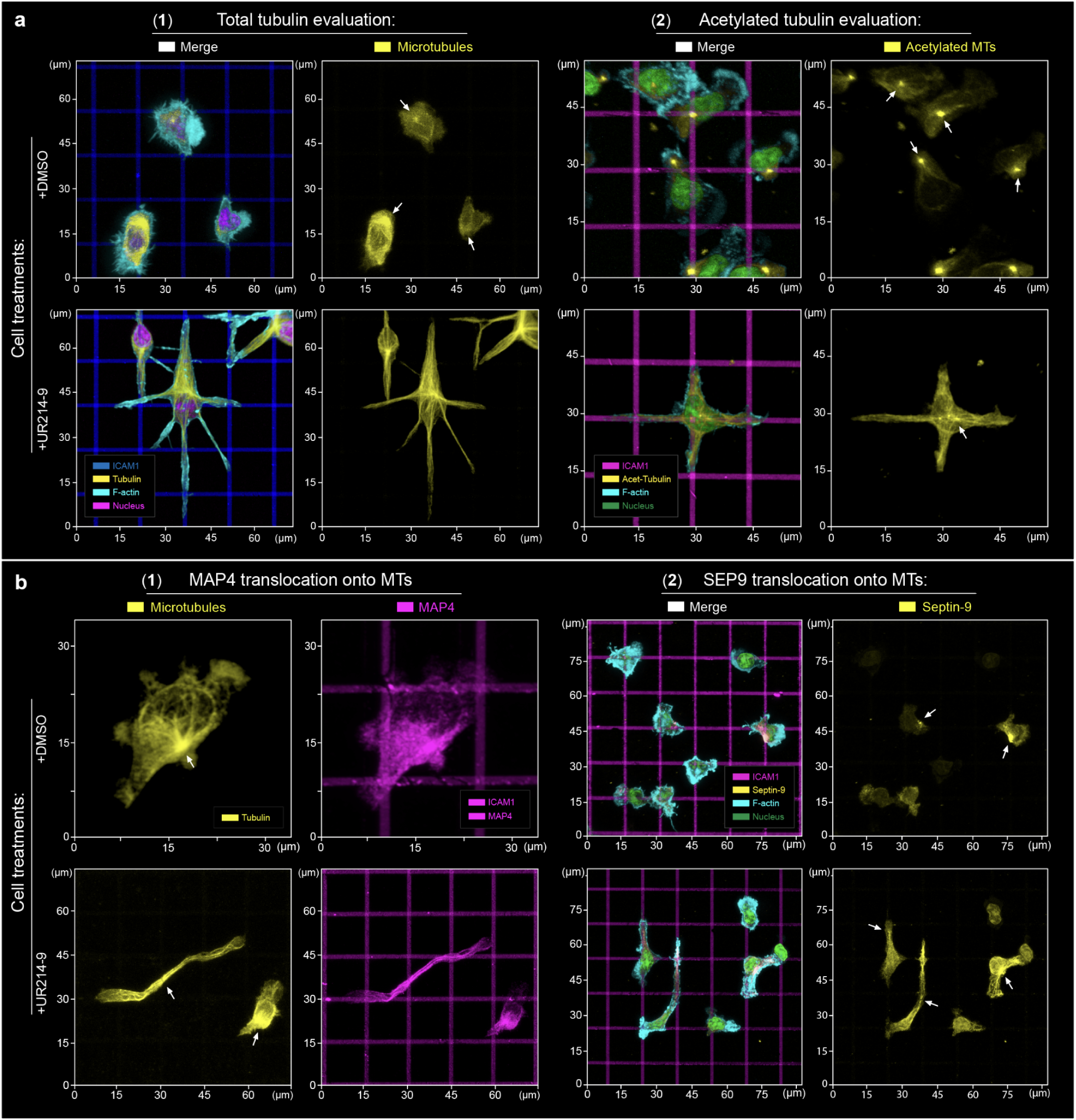
Immunofluorescence analysis of the post-translational microtubule modifications in UR214-9-treated cells. **(a)** Both microtubules density (**1**), and MT acetylation (**2**) are substantially enhanced upon UR214-9-induced suppression of septin GTPase activity **(+UR214-9)**, compared to the diffuse cytoplasmic tubulin and sparse and short microtubules in the control T cells **(+DMSO)**. **(b)** Both MAP4 (**1**) and SEP9 (**2**) translocate to the microtubules upon inhibition of septins with UR214-9.

**Supplemental Figure SI5.**
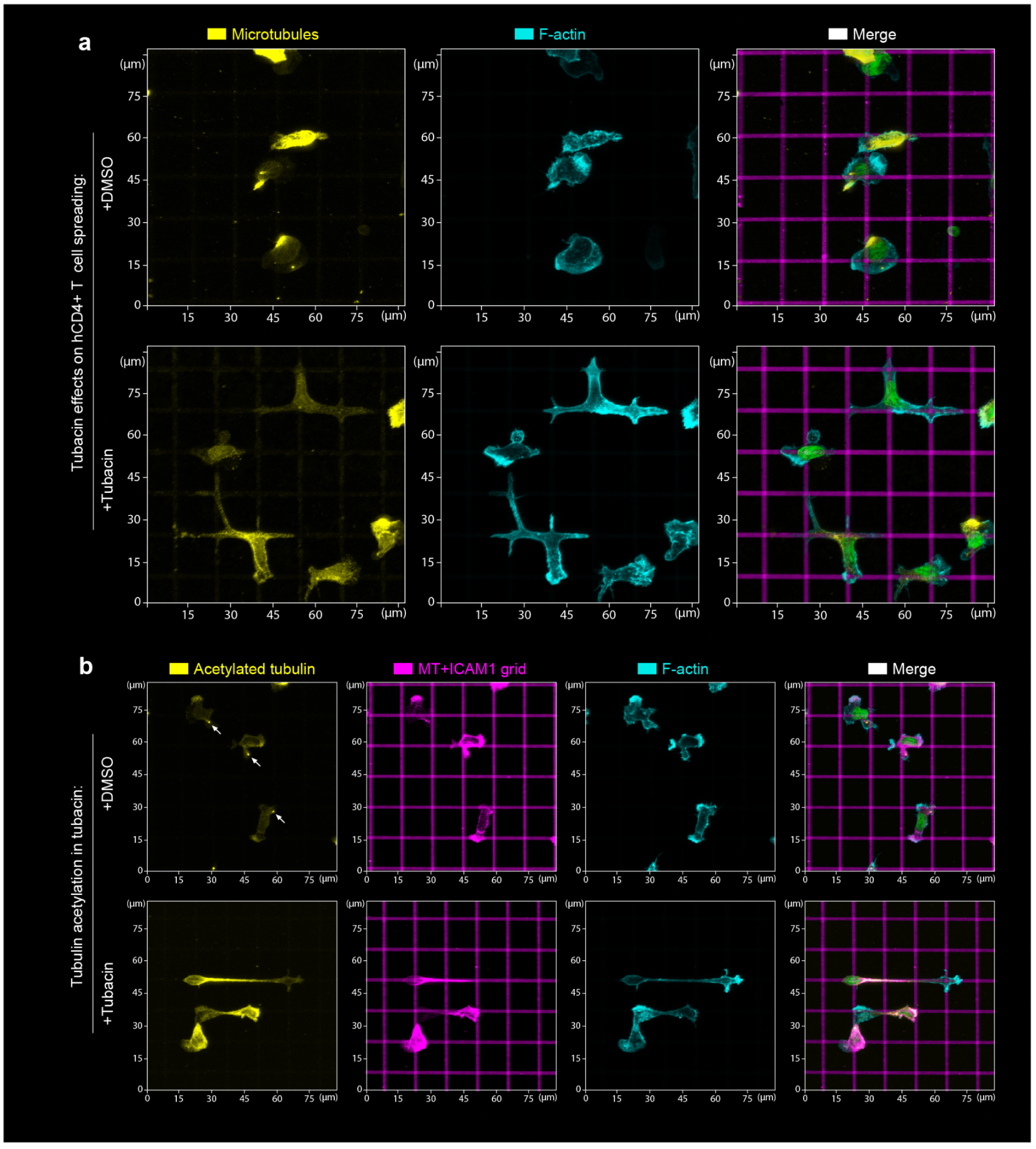
Stabilization of microtubules *via* Tubacin-induced hyperacetylation replicates (*i.e.*, phenocopies) the UR214-9-induced hCD4+ T cell transitioning towards mesenchymal-like dendritic spreading. **(a)** hCD4+ T cells during HDAC6 suppression **(+Tubacin,** t≥1 hour) display a partial amoeboid-to-mesenchymal-like transition, *i.e.* T cell dendritic spreading along the ICAM1 grids. **(b)** Analysis of MT acetylation in hCD4+ T cells, induced by HDAC6 suppression **(+Tubacin,** t≥1 hour), shows an increased acetylation of microtubules compared to the control T cell group **(+DMSO)**. *Identified centrosomes indicated with arrows*.

